# Shortcut barcoding and early pooling for scalable multiplex single-cell reduced-representation CpG methylation sequencing at single nucleotide resolution

**DOI:** 10.1101/2023.05.22.541674

**Authors:** Liyao Mai, Zebin Wen, Yulong Zhang, Yu Gao, Guanchuan Lin, Zhiwei Lian, Xiang Yang, Jingjing Zhou, Xianwei Lin, Chaochao Luo, Wanwan Peng, Caiming Chen, Duolian Liu, Junxiao Zhang, Sadie L. Marjani, Qian Tao, Xuedong Wu, Sherman M. Weissman, Xinghua Pan

**Affiliations:** Department of Biochemistry and Molecular Biology, School of Basic Medical Sciences, Southern Medical University, and Guangdong Provincial Key Laboratory of Single Cell Technology and Application, Guangzhou 510515, Guangdong Province, China; Department of Hepatobiliary Surgery II, Zhujiang Hospital, Southern Medical University, Guangzhou 510280, Guangdong Province, China; SequMed Institute of Biomedical Sciences, Guangzhou 510530, Guangdong Province, China; Department of Biology, Central Connecticut State University, New Britain, CT 06050, USA; Cancer Epigenetics Laboratory, Department of Clinical Oncology, State Key Laboratory of Oncology in South China, Sir YK Pao Center for Cancer and Li Ka Shing Institute of Health Sciences, Chinese University of Hong Kong, 999077, Hong Kong, China; Department of Pediatrics, Nanfang Hospital, Southern Medical University, Guangzhou 510515, Guangdong Province, China; Department of Genetics, Yale School of Medicine, New Haven, CT 06520, USA; Guangdong-Hongkong-Macao Great Bar Area Center for Brain Science and Brain-Inspired Intelligence, Guangzhou 510515, Guangdong Province, China; Shenzhen Bay Laboratory, Shenzhen, 518035, Guangdong, China

**Keywords:** single-cell epigenomics, DNA methylation, bisulfite conversion, CpG Island, scalable throughput, traceable cell identity

## Abstract

DNA methylation is essential for a wide variety of biological processes, yet the development of a highly efficient and robust technology remains a challenge for routine single-cell analysis. We developed a multiplex scalable single-cell reduced representation bisulfite sequencing (msRRBS) technology with off-the-shelf reagents and equipment. It allows cell-specific barcoded DNA fragments of individual cells to be pooled before bisulfite conversion, free of enzymatic modification or physical capture of the DNA ends, and achieves unparalleled read mapping rates of 62.51%, covering 59.95% of CpG islands and 71.62% of promoters in K562 cells on average. Its reproducibility is shown in duplicates of bulk cells with near perfect correlation (R=0.97-99). At a low 1 Mb of clean reads, msRRBS provides consistent coverage of CpG islands and promoters, outperforming the conventional methods with orders of magnitude reduction in cost. Here, we use this method to characterize the distinct methylation patterns and cellular heterogeneity of 6 cell lines, and leukemia and hepatocellular carcinoma models. Taking 4 hours of hands-on time, msRRBS offers a unique, highly efficient approach for dissecting methylation heterogeneity in a variety of multicellular systems.

## INTRODUCTION

Cytosine methylation at DNA CpG dinucleotides, or DNA methylation (DNAme), is an important epigenetic mechanism that regulates gene expression in health and disease. Whole genome bisulfite sequencing (WGBS)^1^ and reduced representation bisulfite sequencing (RRBS)^2^ accurately quantify the methylation of cytosine at single-base resolution genome-scale, and remain the gold standard for CpG methylation analysis. More recently, single-cell CpG methylation sequencing has shown its unique ability to explore the heterogeneity of the epigenetic state among cells in a variety of biological systems^3–7^. In addition, it is highly beneficial for profiling of the methylomes of cells from rare sources, such as mammalian blastomeres and some liquid biopsies, which are difficult to examine with bulk analysis methods^8, 9^. The first single-cell RRBS (scRRBS) report digitally detected the methylation status of single CpG sites in haploid sperm, covering up to 1.5 million CpG sites in the mouse genome^8^. Then, single-cell WGBS, scBS^10^, was developed using post-bisulfite adapter tagging (PBAT)^10–12^, which could detect up to 48.4% (10.1 of 20.87 million) CpG sites within the mouse genome. All of these methods^8, 10, 12^ are based on independent processing of each single cell from cell isolation to library construction with PCR.

Early sample-pooling greatly reduces DNA damage and the cumbersome cell-by-cell bisulfite conversion and post processing. There have been only a few techniques that are able to pool multiple single cells before bisulfite conversion until recently. The outstanding methods for high-throughput single-cell WGBS are represented by snmC-seq2^13^ and sci-METv2^5^, showing their power in the dissection of cellular heterogeneity of the DNA methylome despite the shallow depth of coverage. Although the WGBS strategy theoretically profiles the entire genome at each cytosine, it is cost-prohibitive when a large number of individual cells require analysis and high coverage of CpG sites is desired. Moreover, PBAT, used in scWGBS, prevents the labeling and pooling of every single cell before bisulfite conversion, which results in a practical pattern where only sparse and usually inconsistent CpG sites are detectable across single cells.

In contrast, RRBS enriches the CpG dinucleotide-dense regions that contain key elements in CpG methylation regulation, like CpG islands, promoters, and other important gene regulatory elements (or regions). This makes RRBS a more sequencing-cost efficient method than WGBS, and it allows for deeper coverage of areas of interest, albeit with a low alignment rate. Adapting the RRBS principle^14–16^, we have developed an extraordinary strategy for scalable, multiplex single-cell RRBS (msRRBS). msRRBS enables a panel of single cells to be pooled and bisulfite converted in a single tube, immediately after cell lysis and restriction enzyme digestion, followed by a novel shortcut ligation of barcode-adapters without any enzymatic modification or physical capture of the target DNA fragments. msRRBS is highly efficient in library construction, with consistent coverage of the key elements, as well as orders of magnitude reduction in the cost per cell, while robust results are obtained as long as the genomic integrity of the cell is retained. Furthermore, sequencing data is tracible to each input cell, and the multiplex scale is extendable to high throughput. Our results are based on tests of hundreds of single cells and low-input tissue samples from both humans and mice generated from 8 cell lines, cytarabine-resistant myeloid leukemia cells, and hepatocellular carcinoma cells from a mouse model.

## MATERIALS AND METHODS

### Preparation of samples

Cell lines K562, GM12878, 293T, MGC803, MDA-MB-231, HT22 and CT26 were grown in their suggested media supplemented with 1% penicillin and streptomycin and maintained in a 37°C humidity-controlled incubator with 5.0% CO_2_.

The cytarabine (Ara-C)-resistant KG-1a cell line (KG-1a_R) was generated by direct high-dose drug shock on KG-1a, which is closer to the practical scenario in clinics. Specifically, the KG-1a cell line cultured in vitro with DMEM medium containing 10% fetal bovine serum was treated with 1 μM Ara-C for 3 days, followed by a 9-day drug holiday to restore cell numbers, while the medium was changed every 3 days. Subsequently, KG-1a_R was resistant to 1 μM Ara-C, and 1 μM Ara-C was added to cell culture to maintain the resistance.

Along with three healthy mouse controls, two hepatocarcinoma C57BL/6 mice aged 4-5 months were used, which were generated with liver-specific MYC overexpression, approved by the Southern Medical University Experimental Animal Ethics Committee and conducted in strict accordance with the Committee’s guidelines. DNA was extracted from cell line K562 and liver tissues of HCC mice and healthy mice with the cell/tissue genomic DNA extraction kit (Tiangen, cat. no. DP304) according to the manufacturer’s instructions.

### msRRBS library preparation

Firstly, 2 adapters were generated. For barcode adapter (AdBc), oligonucleotide AdBcS (S = short) four times the number of moles of the AdBcL (L = long) were annealed with 1x T4 DNA Ligase Reaction Buffer (New England Biolabs, Cat. no. B0201S) and heated to 94°C for 3 min, and then rapidly cooled to 80°C, followed by slow cooling to 4°C at a rate of 0.1°C s−1. For sequencing primer adapter AdSp, an equal volume of moles of 2 oligonucleotides was used for the annealing process described above. All adapters are available in supplementary tables 1 and 2.

To digest the genome of each cell, 2.5 µl lysis mixture, containing 0.125µl protease (Qiagen, cat. no. 19155), 0.075μl 10% triton X-100, 10mM Tris-EDTA, 20mM KCl and 2μl nuclease-free water, was added into each well (or tube) with a single cell contained in 1 µl PBS. After incubation at 50°C for 15 min, the wells were heated at 70°C for 30 min. In each of the lysis mixture (3.5 µl) wells, 36.5 µl was added, containing 27.5 µl nuclease-free water, 1 µl carrier RNA (1µg, from Qiagen, cat. no. 59824), 4 µl of Dr.GenTLE precipitation carrier (Takara, cat. no. 9094) and 4 µl sodium acetate. After mixing the contents of the well, 112 µl of pre-cooled 100% ethanol (-20°C) was supplied to each well. The strips of the wells or plates were then incubated at -20°C for 10 min, and centrifuged at 13,300 rpm at 4°C for 15 min. Then, the supernatant in each well was removed carefully to avoid disturbing the white precipitate, and the genomic DNA pellet was washed with 200 µl of 80% ethanol (-20°C) and then the wells were centrifuged at 10,000 rpm at 4°C for 10 min. The residual liquid was discarded, and the genomic DNA in each well was air dried with caps open until its color turned transparent. The genomic DNA was then incubated in a 3 µl mixture that contained 0.1 µl nuclease-free water, 1 µg carrier DNA, 1× Tango buffer (Thermo Fisher, cat. no. BY5), 6 U MspI (Thermo Fisher, cat. no. ER0541, 10 U/µl) and 60 fg unmethylated lambda-DNA (Promega, cat. no. D1521) at 37°C for 2.5 h.

AdBc ligation was performed in 5 µl solution containing the 3 µl of digested DNA from the previous step, 0.6 µM AdBc, 0.2 µl of Fast-Link DNA Ligase (Lucigen, cat. no. LK6201H), 1.2 mM ATP, 0.3 µl 10× fast link ligation buffer and 0.6 µl of nuclease-free water at 25°C for 20 min, followed by incubation at 16°C for 15 min (the temperature can be held up to 14 h) and 20 min at 25°C, followed by 75°C for 20 min. Finally, 25 mM EDTA (Thermo Fisher, cat. no. AM9260G) was added to the tubes, which were then incubated at 37°C for 15 min with the lid set at 50°C; after incubation the samples were spun down. Each batch of processed samples was pooled in a new tube, 200 µl or 1.5 ml, according to sample number. The pooled DNA was purified using 1.5× volume of AMPure XP beads (Beckman coulter, cat. no. A63880) following the manufacturer’s instructions, and a total of 18 µl DNA solution was collected. Then, 2 µl adapter filling mixture containing 0.375 mM 5-Methylcytosine dNTP Mix (Zymo Research, cat. no. D1030), 0.75 µl of Thermopol Reaction buffer (New England Biolabs, cat. no. B9004S, 10×) and 0.5 µl of Sulfolobus DNA polymerase IV (New England Biolabs, cat. no. M0327S) was added to the 18 µl DNA solution and incubated at 55°C for 30 min.

The bisulfite conversion mixture was made of 1 µg carrier DNA, 19 µl nuclease-free water, 85 µl bisulfite solution and 15 µl of protect buffer (Qiagen, cat. no. 59824). The volume of 20 µl of DNA solution (after filling) was incubated in 120 µl bisulfite conversion mixture at 95°C for 5 min, 60°C for 10 min, 95°C for 5 min and 60°C for 20 min. Purification was performed according to manufacturer’s instructions and the converted DNA was then eluted in 32.5 µl nuclease-free water (17 µl × 2 times) through a column from the DNA Clean & Concentrator kit (Zymo Research, cat. no. D4003).

To generate the final library, a 50 µl mixture for the first-round of PCR was prepared with 32.5 µl bisulfite converted DNA solution, 2.5 U Takara Epi Taq HS (Takara, cat. no. R110A, 5 U/µl), 2.5 mM MgCl_2_, 5 µl 10× Takara Taq PCR buffer (Mg^2+^ Free), 0.1 mM dNTP and 1 µM PreAP. Thermal cycling reaction was performed as follows: 95°C for 5 min; 25 cycles of (95°C 30 s, 56°C 30 s, 72°C 45 s); 72°C for 10 min; 4°C hold. Next, the PCR products were purified via DNA Clean & Concentrator kit (Zymo Research, cat. no. D4003), and eluted in 17 µl of nuclease-free water. This 17 µl of DNA solution was incubated with 3 µl digestion mixture containing 1 µl BciVI (New England Biolabs, cat. no. R0596S) and 2 µl CutSmart buffer (New England Biolabs, cat. no. B7204S, 10×) at 37°C for 2h, followed by heating at 65°C for 20 min. After that, purification was performed using the DNA Clean & Concentrator kit and the DNA was eluted with 15 µl nuclease-free water. A 25 µl ligation mixture was prepared by adding the 15 µl DNA solution from the previous step, 5 µM sequencing adapter AdBc, 1µl Fast-Link DNA ligase (Lucigen, cat. no. LK6201H), 3 µl 10× Fast link ligation buffer and 1mM ATP. Ligation reaction and inactivation of enzymes were performed as described above. Finally, a total of 20 µl DNA solution was collected using DNA Clean & Concentrator kit. DNA fragments between 125-300 bp (size extendable up to 700 bp) were recovered by 2% E-gel (Invitrogen, cat. no. G402002) and purified using Zymo Gel DNA recovery kit (Zymo Research, cat. no. D4007) following the manufacturer’s instructions. A Qubit fluorometer (Thermo Fisher, cat. no. Q33238) was used to measure the concentration of the eluted DNA solution. The PCR mixture was prepared with approximately 10 ng of DNA recovered by size selection on gel, 12.5 µl GC-rich Phusion High-fidelity 2× master mix (Thermo Fisher, cat. no. F532S), 1 µM primer P5, 1 µM primer P7 and nuclease free water up to 25 μl final volume. A final library amplification was carried out with the program: 95°C for 1 min; 4-5 cycles of (95°C 30 s, 57°C 30 s, 72°C 45 s); 72°C for 10 min; 4°C hold. The DNA Clean & Concentrator kit was used to clean up excess library primers, and 20 µl of the eluted library was used for the final size selection, as descried above. Finally, a total of 25 µl of DNA fragments of 175-350 bp in length (the upper limit is extendable to 550 or 750 bp when specified) was obtained. Optimal concentration of the DNA solution from the second round of DNA size selection was >= 2 ng/µl. All libraries were sequenced using the Illumina HiSeq X Ten platform with 10-30% PhiX spike-in and 150-bp paired-end reads.

### Data analysis

The 4-bp of marker sequence, GGTG, or a random sequence, NNNN, and 6-bp sample barcode were located at the beginning of both reads. FASTQ files were first split by pre-assigned sample barcodes. No mismatches were allowed. If the average sequencing quality of the barcode was lower than Q20, the whole read was filtered out. The marker sequence, sample barcode and Illumina adapters were trimmed from read 1 and read 2 with Cutadapt^17^ and Trimmomatic^18^. Notably, the 3’ end of the reads often contained the inverse complementary sequence of the fixed sequence and sample barcode because the recovered fragments of the library were approximately 30-200 bp if it was not specified (otherwise, 30-400 bp or 30-600 bp), and the sequencing strategy was 2 × 150 bp.

Clean reads were then aligned to the reference genome (GRCh38 or GRCm38) in paired-end mode using the Bismark^19^ tool. Calculation of the conversion rate and DNAme calling were performed as descried^20^. CGmapTools^21^ software package was used to annotate the genome region of CpG sites, so as to obtain the corresponding information of different CpG sites on chromosomes. Regions include CpG islands, CGI shores, CGI shelves, promoters (containing CGIs and 1500 bp upstream and 500 bp downstream of all transcription start sites of protein-coding genes), genes, exons, transcripts, CTCF binding sites and enhancers downloaded from the UCSC Genome Browser. Duplication or redundant reads of genomic regions were removed according to the coordinates of the chromosomes, that is, only one of the genomic regions with identical genomic coordinates remained.

A methylation signal matrix was used to calculate the correlation between different samples or groups. Cluster analysis through principal component analysis (PCA), t-distributed Stochastic Neighbor Embedding (tSNE), and multidimensional scaling (MDS) was also performed. CalculateDiffMeth^22^ function or DSS package was used for differential analysis of CpG sites or elements (DMC or DMR), and the obtained results were visualized by heat map or volcano map to display the differential DNAme information between different samples or groups. Homer^23^ was used to annotate the DMRs according to reference genomic elements to explore the potential phenotypes and functions corresponding to differential methylation levels in different samples. Differentially methylated genes were screened for GO or KEGG pathway enrichment to reveal the potential relationship between methylation-level information and phenotype.

## RESULTS

### msRRBS procedure overview

We sought to provide a middle to high-throughput RRBS protocol (**Figure 1 and Supplementary Figure 1a**) through early labeling of each cell for multiple single cells in parallel, followed by subsequent pooling of the panel of cells for single-tube transformation. Thus, it makes the procedure highly efficient and robust in operation with lower cost, while generating high quality data consistent among the cells. A summary of the protocol follows. First, a panel of single cells in different wells or tubes obtained by pipette aspiration, flow sorting, or microfluidic delivery are lysed to fully expose genomic DNA. Second, CpG-rich DNA regions of the genome for each cell are fragmented with the MspI enzyme or its alternatives. Third, the double-stranded genomic fragments with sticky ends from each cell are directly (without end repairing and A-tailing) ligated to a specifically designed adapter (AdBc), composed of a longer strand and a shorter strand (AdBcL and AdBcS). AdBcL is built with the cell-barcode at its 3’ end consisting of a combination of 6 nucleotides to label each cell, and the BciVI recognition site (5’-GTATCC-3’) follows the barcode; its 5’ end is built with a tag for priming PCR amplification after bisulfite conversion, in which all the cytosines (Cs) in the AdBcL are methylated. AdBcS is complementary to the 3’ end of AdBcL. After ligation, the ligase is inactivated by heating, and AdBcS, whose 5’ end is not phosphorylated and not covalently bound to the 3’ end of the digested DNA fragments, is dissociated away, while AdBcL keeps its covalent connection with the genomic fragments. Fourth, all constructs of adapter-DNA fragments barcoded for each cell for the panel of cells, in which each is built with different barcode, are then pooled into a single reaction tube for subsequent processes. Fifth, Sulfolobus DNA polymerase IV is employed to fill in the adapter by an extension of the 3’ end of the DNA fragments along AdBcL; the reaction includes methylated C but no unmethylated C. Sixth, bisulfite is used to convert unmethylated C to U. Seventh, the first round of PCR is performed with a limited number of cycles. Eighth, the adapter/primer portion of the amplicon is excised with BciVI, and the barcodes of the cell are retained. Importantly, this cleavage leaves an ’A’ tail at the 3’ end of the dsDNA to enable the ligation of the conventional library adapter that has a ’T’ tail at its 3’ end for the second round of PCR (**Supplementary Figure 1b**). Tenth, a second round of PCR amplification is performed incorporating an index for each pool of cells; library products of 175-350 bp (alternatively 550 bp or 750 bp as the upper limits when specified) are then isolated by size selection through agarose gel or its alternatives. After purification, the obtained library is sequenced (**Supplementary Figure 1c**). An aliquot of purified genomic DNA or a small amount (micro-bulk) of cells can be alternatively processed as a sample instead of a single cell.

**Figure 1.**
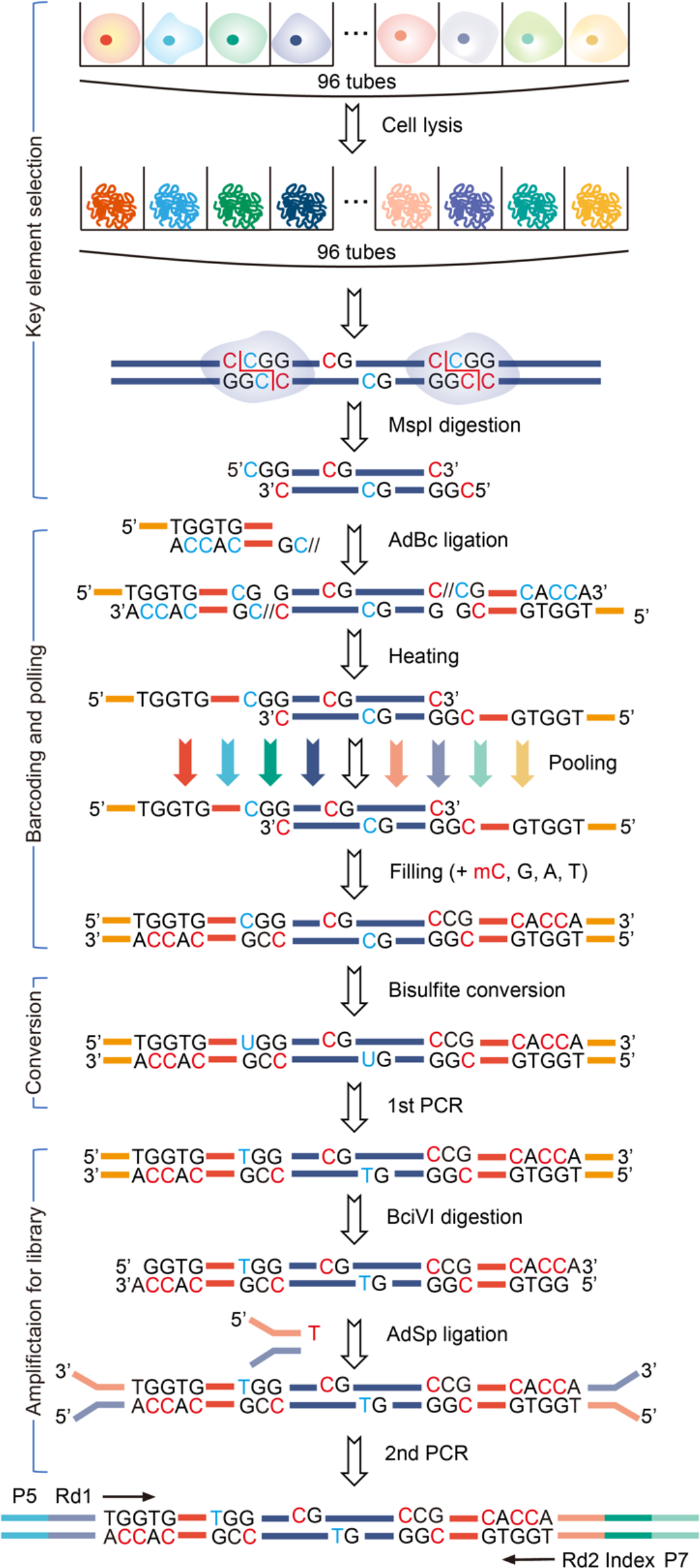
Procedure scheme of msRRBS. After lysis and Msp I digestion of multiple single cells in parallel independently, genomic DNA fragments of each single cell are directly ligated to a DNA adapter (AdBc) with a cell-specific barcode composed of a relatively long strand (AdBcL) and a short strand (AdBcS). Because AdBcS is non-covalently held at its 5’ end to the genomic DNA fragment at 3’ end, it falls off the AdBcL when the ligase is inactivated at 75°C. Then multiple single cells are pooled into a single tube. With addition of 4 nucleotides including methylated C, while no unmethylated C is included, the new AdBcS is re-synthesized along AdBcL with a DNA polymerase, Sulfolobus DNA polymerase IV, to re-construct the double strand adapter. Here bisulfite conversion is executed, and the products are recovered by PCR amplification. The amplicon is digested with BciVI, and a sticky end with an ‘A’ tail protruding at 3’ end is generated, which enables this product to be directly ligated to a conventional Y-shaped adapter with Illumina P5 and P7 holding a ‘T’ tail protruding at 3’ end. Finally, a minimum cycle number of PCR is followed to finalize the sequencing library.

### Improved mapping rate and coverage of single cells achieved by msRRBS

We analyzed the basic features of msRRBS using the cell line K562. The average conversion rate was obtained by spiking 60 fg of unmethylated lambda DNA in the pooled samples for the msRRBS assay, which was 99.64% (ranging from 99.54% to 99.86%), indicating that unmethylated Cs were efficiently converted to Ts in both single cell and bulk samples (**Supplementary table 3 and 4**). Therefore, the methylation status of CpG sites obtained in the sample data is reliable. Indeed, the average DNAme levels of cytosines in the CHG and CHH context, which are known to be nearly unmethylated in most mammalian cell types, were detected as being less than 0.43%, further confirming the complete conversion.

We also found that the increased sizes of target fragments in the sequencing library did not significantly raise the mapping rate. When the library sizes were recovered as 175-350 bp, 350-550 bp, 175-550 bp and 175-750 bp, the average mapping rates of raw reads in single cells (msRRBS) were 57.26% ± 3.65%, 58.83% ± 0.59%, 58.10% ± 1.19% and 62.51% ± 3.85%, respectively. In bulk cells (mRRBS), the average mapping rates were higher than that of single cells. However, the long range of fragments sequenced in bulk cells overall did not significantly result in higher mapping rates than the short range of fragments (**Figure 2a)**. For individual samples, the mapping rate was up to 79.85% in single cells and to 89.51% in bulk cells over the whole study.

**Figure 2.**
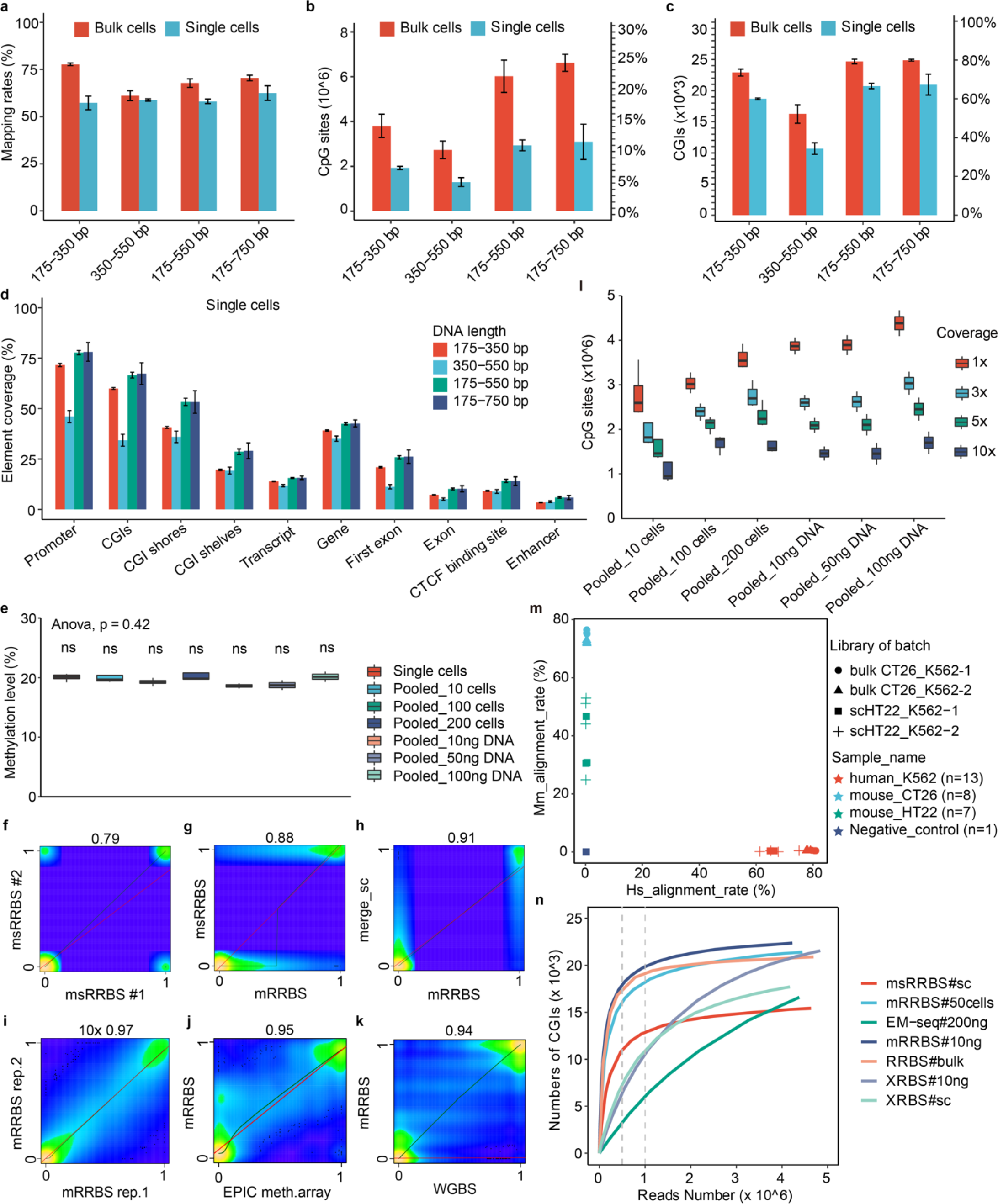
Basic features of msRRBS. The mapping rate (a), detection rate of CpG sites (b) and CGIs (c) associated with different library sizes in single (n=3-4) and bulk cells (n=4). (d) Coverage of functional elements or regions along the genome associated with different library sizes, detected in K562 single cells (n=3-4). (e) Overall methylation level among single cells (n=15), 10-200 cells (n=4) and 10-200 ng DNA (n=2) extracted from K562 cells (p value = 0.42). (f-i) Pearson correlation compares methylation profiles of CpG sites acquired by msRRBS between two single cells (f, R = 0.79), a single cell and a bulk of cells (g, R = 0.88), merged single cells and bulk cells (h, R = 0.91), technical replicates of bulk cells (i, coverage = 10×, R = 0.97) for cell line K562, mRRBS and EPIC methylation array (j, R = 0.95) for KG-1a cells, and public WGBS (k, R = 0.94) for K562 cells. (l) Number of CpG sites detected in micro-bulk (10-200 cells, n=4) and extracted gDNA (10-100ng, n=2) of K562 with different depths of coverage. (m) Mapping rates that align specifically to the mouse genome (x axis) versus the human genome (y axis), confirming no detectable cross-contamination between human and mouse cells prepared in the same reaction pool. Each dot represents a sample (a bulk or a single cell). Duplicate pools of CT26 and K562 bulk cells (∼100 cells), and duplicate pools of HT22 and K562 single cell were independently tested. (n) Number of CGIs with at least 1 CpG site as a function of sequencing depth for msRRBS, mRRBS, conventional RRBS, EM-Seq, and XRBS, detected in K562. The vertical gray dashed lines indicate 0.5 M and 1 M clean reads serving as a threshold for the comparison.

Importantly, when library size range increased from 175-350 bp to 175-750 bp, the coverage was raised accordingly. For the two ranges of library size, the best detection values were respectively up to 2,196,284 and 3,791,693 CpG sites, and up to 20,312 and 22,383 CGIs (corresponding to 72.54% and 79.94% CGIs) in single-cell samples. These values were 4,673,052 and 7,188,812 CpG sites, and 24,051 and 25,115 CGIs (corresponding to 85.89% and 89.70% CGIs) in bulk samples.

When the library size ranges were between 175-350 bp, 350-550 bp, 175-550 bp (generated by silicon-merging the data of 175-350 bp and 350-550 bp) and 175-750 bp, the average detection rates of CpG sites in single cells were 1.93 million, 1.30 million, 2.94 million and 3.09 million, respectively. While in bulk cells, these average detection rates were correspondingly 3.81 million, 2.74 million, 6.08 million and 6.62 million **(Figure 2b**). Compared with 175-350 bp, the detection of CpG sites of 175-550 bp libraries increased by 37.34%. Interestingly the CpG sites detected in the 175-750 bp libraries did not significantly outperform the 175-550 bp libraries. Similarly, the coverage of the functional elements, particularly CGIs **(Figure 2c**) and promoters (**Supplementary Figure 2a**), did not show any noticeably extension when the library sizes were increased. Obviously, the 175-550 bp library results in more CpG sites detected in each functional element, i.e., CGI and promoter, than 175-350 bp library, which should improve the measurement representation of the elements.

Accordingly, this method detected more elements in bulk samples (mRRBS) than in single cells (msRRBS): promoter (81.61% ± 2.52% vs. 71.62% ± 1.58%) and CpG islands (73.52% ± 5.56% vs. 59.95% ± 1.44%) for 175-350 bp library, and a similar trend was noted for wider ranges of library size. Other elements were also covered (**Figure 2c and 2d, Supplementary Figure 2b, Supplementary table 4**).

### Outstanding reliability, reproducibility and efficiency enabled by msRRBS

As most CpG sites are enriched in short fragments with the 4-nucleotide restriction endonuclease MspI, for the ultimate sake of high throughput msRRBS with a minimal sequencing cost, we focused on the library sizes of 175-350 bp for the following analyses. At first, we tested 15 high-quality single cells in a batch as well as its corresponding bulk cells. For single cells, micro-bulk quantities of 10 - 200 cells and 10 - 200 ng of DNA extracted from K562 cells, there was no significant difference in CpG site methylation levels between groups (**Figure 2e**, P > 0.05). It was found that the correlation coefficient (R) of the methylation profiles of two typical single K562 cells was 0.79 (**Figure 2f**), while this factor did reach up to 0.88 (**Figure 2g**) when the correlation analysis was calculated between a single cell and a micro-bulk sample of approximately 50 cells. In addition, the correlation (R) reached 0.91 between the merged single cells analyzed by the msRRBS procedure, and true-bulk cells (**Figure 2h**). Considering the heterogeneity among single cells, we randomly merged the data of 4 single K562 cell samples and identified that the methylation pattern of CpG islands (**Supplementary Figure 2c**) and promoters (**Supplementary Figure 2d**) among cells were also basically similar.

To further assess the reproducibility of msRRBS and rule out the effect of heterogeneity among single cells, the correlation between the two bulk samples (50ng) of K562 cells was measured, with 0.97 (R) at coverage ≥10×(**Figure 2i**). When analyzing CpG sites at depths of ≥3×, ≥ ×, ≥15× and ≥20×, the correlation coefficients (R) were 0.93 (**Supplementary Figure 2e**), 0.95 (**Supplementary Figure 2f**), 0.97 (**Supplementary Figure 2g**) and 0.98 (**Supplementary Figure 2h**), respectively, indicating high robustness of the method. The overall methylation pattern of the co-detected CpG sites by mRRBS was highly consistent with that obtained by EPIC array analysis (**Figure 2j,** R = 0.95) of KG-1a, and a publicly available WGBS dataset of K562 (**Figure 2k,** R = 0.94).

To address the coverage limitation and methylation level accuracy with different amounts of input, we compared micro-bulk samples versus single cells and a very low number of cells. When 200 K562 cells were pooled together, there were no significant differences in detection rate of CpG sites compared with pooled 100 cells, 10ng, 50ng and 100ng DNA (p > 0.05, t-test), but different when compared with 10 cells (p < 0.01, t-test) **(Figure 2l**). As expected, single-cell and bulk-cell samples displayed a consistent methylation level for each functional element over the genome (**Supplementary Figure 2i**).

To evaluate any possible cross-contamination between cells or barcodes, 4 single cells of cell line HT22 (mouse), 4 single cells of K562 (human) and 1 sample of nuclease-free water as a negative control were pooled into the same conversion reaction and library amplification steps. In addition, 4 aliquots of 100 cells of CT26 (mouse) and 4 aliquots of 100 cells of K562 (human) were pooled into the same reaction tube. Two batches of each mix were performed. Mapping of msRRBS data to the genomes from both species confirmed that, technically, there was no evident barcode cross-contamination (**Figure 2m**). Notably, the mapping rate of the negative control to both human and mouse genomes was less than 1%.

With different sequencing depths, it was also recognized that msRRBS reached plateau earlier (0.5-1M), in both bulk-cell and single-cell samples, than when compared to other competing methods, such as EM-seq^24^, which is an alternative WGBS that enzymatically converts unmethylated C genome-wide, and XRBS^14^, an extended scRRBS method. msRRBS, at a relatively low sequencing depth, also outperforms WGBS and scRRBS in overall coverage and consistency (**Figure 2n and Supplementary Figure 2j**). This is attractive for sequencing a large number of single cells for pair-wise comparison and clustering analysis. This result may be attributed to the fact that msRRBS more specifically enriched the regions rich in CG sites, such as CpG islands and promoters. Thus, the coverage of the key epigenetic regulatory elements in msRRBS is more consistent among cells. At a low depth, for example at 0.5 MB reads, msRRBS obtained approximately 1.5 times the CGIs, and 1.3 times the promoters than obtained by XRBS. While bulk cells (mRRBS) had an even higher efficiency than single cells (msRRBS) when compared to XRBS for the coverage of these regions, i.e., approximately 3.5 times the CGIs, and 3 times the promoters than obtained by XRBS. At 1 Mb clean reads, msRRBS still outperformed XRBS. With further, deeper sequencing of single cells, XRBS detected more CpG sites and elements, but msRRBS/mRRBS covered increased CpG sites when sequencing a library with increased range from 170-350 bp to 170-550bp, as demonstrated above. Interestingly, when the sequencing depth is less than 4-5 Mb, bulk-cell sequencing with mRRBS also detects more CpG sites than XRBS. Therefore, msRRBS with 175-350 bp library size is suitable for sequencing a large number of single cells with less sequencing depth, while when more CpG sites are desired, particularly when analyzing rare cells, the 175-550 bp library is preferred.

### msRRBS profiling of methylation characteristics of single cells from distinct cell lines

The relationship of CpG methylation profiles was assayed for 6 cell lines: GM12878, MGC803, MDA-MB-231, 293T, KG-1a and K562, using msRRBS methylation level as the parameter to build correlation maps. We calculated the successful rate of the msRRBS procedure in several batches of the experiment: 86 cells (≥ 0.2 M CpG sites were detected) out of 101 cells tested were used in the following analysis. As expected, 6 different clusters (obtained using tSNE with genomic 0.2-kb windows) were generated with each cell clustered to its corresponding cell line **(Figure 3a)**, and a high correlation was observed among the single cells with each cell line **(Figure 3b, on average R = 0.8)**. We found that 0.2-kb windows best reflect the correlation of the 6 cell lines, followed by three functional elements: CGIs, promoters, and gene bodies **(Supplementary Figure 3a and b)**.

**Figure 3.**
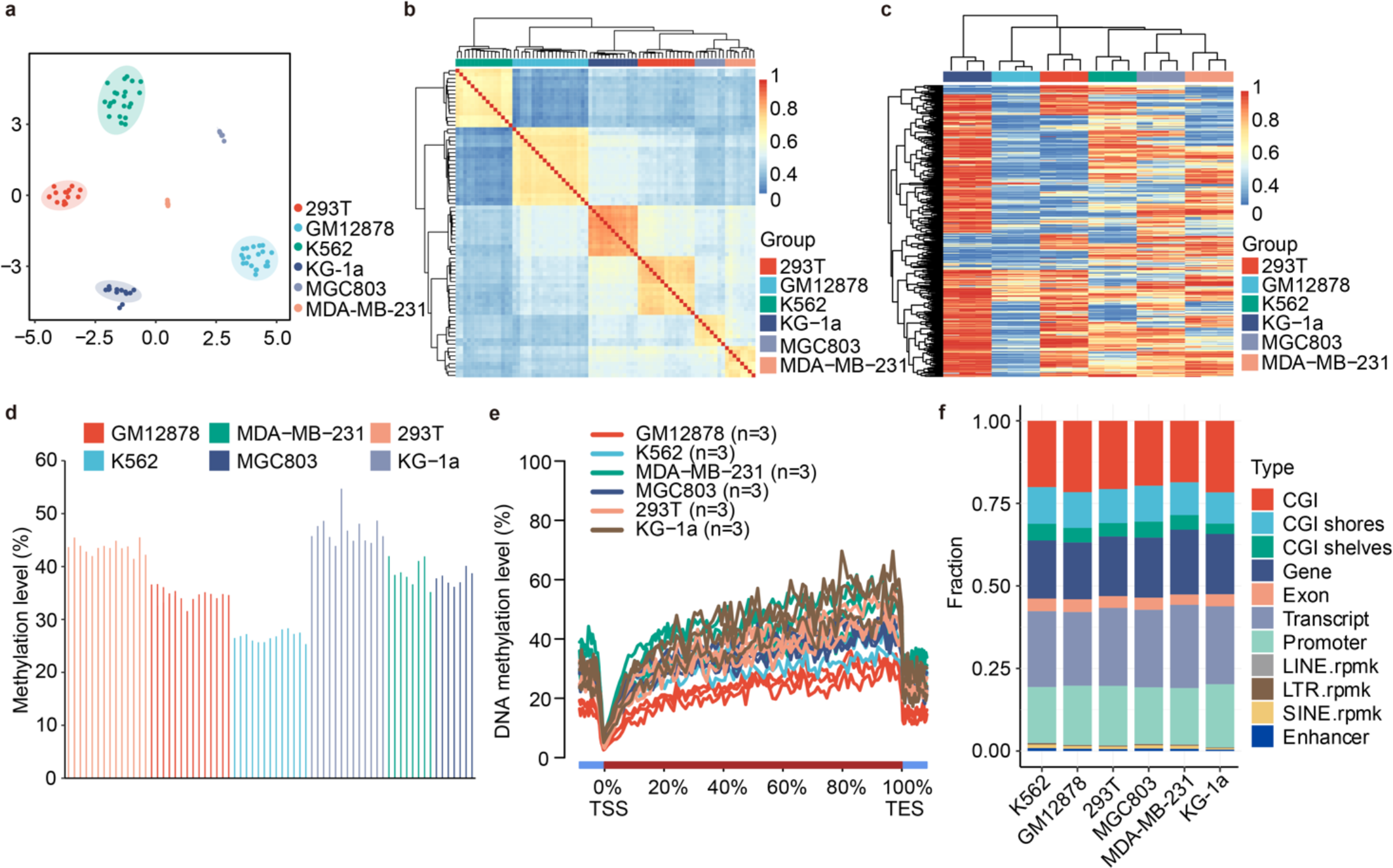
Feasibility and reproducibility of msRRBS profiling for methylation characteristics of a panel of cell lines. (a-b) Single cells from the six cell lines analyzed in 0.2-kb windows over the genome. (a) Unsupervised clustering with tSNE. Six cell lines: 293T (n=15), GM12878 (n=20), K562 (n=22), KG-1a (n=13), MDA-MB-231 (n=8) and MGC803 (n=8). (b) Pearson correlation heatmap. The color key from blue to red indicates low to high correlation. (c) Methylation profile of CGIs for 2, 4, and 8 cells randomly in silico-merged from each of the six cell lines. The number 0 represents unmethylated, and the number 1 represents fully methylated. The color changes from blue to red to indicate a gradual increase in methylation levels. (d) Bar plot shows the average genome-wide DNAme level of CpG sites for the six cell lines. (e) Average DNAme levels along the gene body and 5 kb upstream and downstream of the TSSs and the TESs for all RefSeq genes of the six cell lines. (f) Average detection rates of various genomic regions in each of the six cell lines.

Furthermore, the methylation characteristics of the three elements were examined using randomly merged data of 2, 4 and 8 single cells. The average correlation coefficients (R) of methylation levels based on these 3 elements in the 6 cell lines were 0.97, 0.95 and 0.91, respectively (**Supplementary Figure 3c-e**). In addition, each of the 6 cell lines were distinguished and formed isolated clusters based on methylation profile of CGIs, with obvious distinct patterns among the cell lines (**Figure 3c**); a similar pattern was observed with promoters (**Supplementary Figure 3f**). At the same time, the methylation profile of single cells from the same cell line was relatively consistent with existence of heterogeneity among cells. Average DNAme levels were similar across single cells of a given cell line (34.60 ± 1.28%, 26.75 ± 0.93%, 39.01 ± 2.48%, 37.86 ± 1.32%, 43.57 ± 1.20% and 46.90 ± 2.84 for GM12878, K562, MDA-MB-231, MGC803, 293T and KG-1a, respectively (**Figure 3d**).

The methylation pattern of particular genomic regions was also analyzed with the msRRBS data. Three single cells of each cell line were randomly selected as representatives to draw cross-gene methylation profiles (**Figure 3e**); the results were consistent with early reports^20^. The methylation level of genes across the genome was higher than that in the adjacent regions. There was an expected hypomethylation valley at the transcription start site (TSSs) and methylation levels increased gradually from the 5’ end (TSS) of the gene to the 3’ end (transcriptional end site, or TES). The same result was obtained when 86 cells from 6 cell lines were plotted (**Supplementary Figure 4a**). Meanwhile, we also mapped the methylation levels of CpG islands over the whole genome and revealed that the methylation levels of CpG islands were lower than that of adjacent genomic regions, as shown in the randomly plotted data of 3 single cells or with the whole panel of 86 single cells (**Supplementary Figure 4b and c**). In addition, the promoter regions (**Supplementary Figure 4d**), the first exons (**Supplementary Figure 4e**) and a combination of all exons (**Supplementary Figure 4f**) demonstrated consistent hypomethylation patterns within each type of element among different cell lines. The average number of the eleven functional elements detected in each single cell from the six cell lines remained roughly consistent (**Figure 3f**).

Next, we mapped the entire genome (**Supplementary Figure 4g, left**) and chromosome 1 (**Supplementary Figure 4g, right**) for the msRRBS status for 6 cell lines each with 8 single cells merged. The cell line KG-1a was derived from acute myeloid leukemia and displayed high methylation levels. Both maps revealed that the methylation level of KG-1a was the highest of the 6 cell lines, at about 55%, while K562 had the lowest methylation. Subsequently, corresponding heatmaps of methylation status genome-wide were generated (**Supplementary Figure 4h**), which were consistent with those above. In the region 1,664,226 to 1,664,541 of chromosome 7, a representative lollipop map illustrates the different levels of methylation in the 6 cell lines (for each, data from a single cell was shown), and each cell line displayed distinct features of its own (**Supplementary Figure 4i**).

### Single-cell methylation profiling by msRRBS of cytosine arabinoside (Ara-C) resistant versus the original myeloid leukemia cell line KG-1a

To evaluate the performance of the method in a practical biological system, msRRBS was applied to Ara-C resistant KG-1a cells (KG-1a_R). Thirty-two single-cell libraries, 16 each for KG-1a_R and KG-1a, in addition to the corresponding bulk-cell libraries, were constructed. We filtered out the data for 4 unqualified cells and used data from 28 cells for the downstream analysis, which covered more than 0.2 million CpG sites per genome. A correlation analysis determined that the correlation coefficient (R) between two single cells from same KG-1a_R group was up to 0.87 (**Figure 4a**). The correlation coefficients between a single cell and its corresponding bulk-sample, between the pseudo-bulk (14 cells merged in-silico) and its corresponding bulk, and between biological replicates of bulk samples, were 0.93 (**Figure 4b**), 0.93 (**Figure 4c**), and 0.99 (**Figure 4d**), respectively. The correlation results with the cells of KG-1a were similar (**Supplementary Figure 5a-d**). Two distinct clusters were shown when unsupervised clustering was performed based on the methylation level of either CGIs (**Figure 4e**) or promoters (**Supplementary Figure 5e**) for KG-1a_R (n=11) and KG-1a (n=12). We further generated the methylation curve of gene bodies, promoters, and CGIs separately over the genome for single cells (**Figure 4f**) and bulk cells (**Supplementary Figure 5f**) from the KG-1a versus its counterpart KG-1a_R. Overall, there was no significant difference in methylation levels between these two groups, either in single cells or bulk cells. However, this analysis identified that the fluctuation of the methylation signal for single cells was greater than that of bulk-cell samples in the three elements or regions, CGIs, promoters and gene bodies, probably reflecting the heterogeneity among cells and the limited coverage of CpG sites.

**Figure 4.**
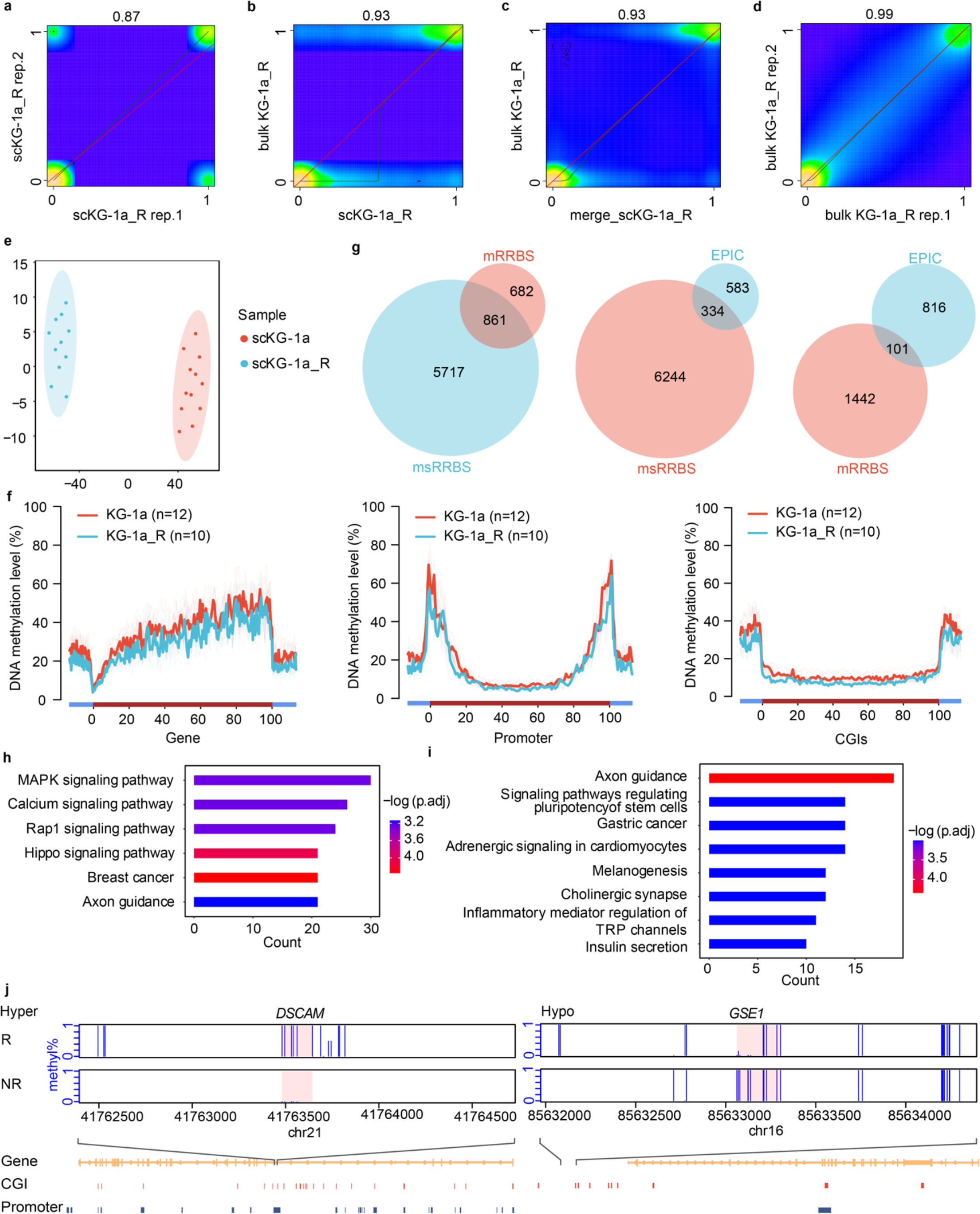
Epigenomic and biological significance of a myeloid leukemia cell culture resistant to cytarabine revealed by msRRBS. (a-d) Pearson correlation heatmap compares methylation values of individual CpG sites acquired by msRRBS between two single cells (a, R = 0.87), between a single cell and a bulk of cells (b, R = 0.93), between pseudo-bulk (merged 14 cells) and bulk cells (c, R = 0.93), and between technical replicates from a bulk of cells (d, R = 0.99) for cytarabine-resistant KG-1a (KG-1a_R). (e) Unsupervised clustering based on single-cell CpG island methylation for KG-1a_R (n=11) and original KG-1a (n=12) with tSNE. (f) Average DNAme patterns across gene body, promoters, and CGIs from the data set of single cells by msRRBS. (g) Hyper-DMR intersection of KG-1a_R vs. KG-1a among the methods msRRBS, mRRBS, and EPIC methylation array. Hyper (h) and hypo (i) DMRs enriched in the KEGG pathway of a bulk of cells are showed for KG-1a_R versus original KG-1a. (j) Two windows each contain a DMR of merged KG-1a single cells from KG-1a_R versus the corresponding original KG-1a population (MWU-test p-value < 0.05, number of CpG sites in one DMR ≥ 3, the distance between two independent DMRs ≥ 100bp, length of DMR ≥ 50bp).

Differential methylation analyses for DMCs and DMRs were performed on the merged single cells **(Supplementary Figure 5g)** and bulk cells (**Supplementary Figure 5h**). We obtained the differential hyper-methylated (**Figure 4g**) and hypo-methylated (**Supplementary Figure 5i**) regions for KG-1a_R versus KG-1a, using msRRBS (merged data of 11 or 12 single cells) and mRRBS (bulk-cell data) compared to the bulk-cell EPIC array, and we calculated the overlap rate between the two methods. We illustrated that merged msRRBS covered more DMRs than the mRRBS bulk results; that merged msRRBS covered more DMRs than the bulk EPIC results; and that bulk mRRBS covered more DMRs than the bulk EPIC results, significantly.

The DMRs obtained above were inspected for functional significance by KEGG analysis using DMRs obtained from bulk cells of KG-1a_R versus KG-1a. We identified enrichment of the MAPK signaling pathway^25^, Calcium signaling pathway^26^, AXON guidance^27^ among others (**Hyper-Figure 4h, hypo-Figure4i**), which have been associated with the occurrence and development of myeloid leukemia. These KEGG pathways that were enriched specifically in DMRs for KG-1a in the bulk samples overlapped in the corresponding KEGG list from single cells. As representative results, the regulatory elements for *DSCAM* and *SBF2* were hyper-methylated, while *GSE1* and *GCNT2* were hypo-methylated, in KG-1a_R in contrast to KG-1a (**Figure 4j and Supplementary Figure 5j**). The role of these genes in leukemia or other cancers is consistent with our result. Specifically, *GCNT2* is overexpressed in highly metastatic breast cancer cell lines of human and mouse origin and breast tumor samples^28^. *GSE1* employs protumorigenic activity in AML cells and is a down-regulation reliever of the differentiation block in AML blast cells^29^. The change of *DSCAM* is a risk factor for Down Syndrome leukemia^30^. Finally, *SBF2-AS1* is significantly upregulated in NSCLC compared with corresponding non-tumor tissues, and a high expression level of *SBF2-AS1* is correlated with lymph node metastasis and the advanced TNM stage of NSCLE^31^.

### Representative methylation pattern of mouse hepatocellular carcinoma (HCC) revealed by mRRBS

Recognizing that msRRBS was highly efficient in bulk-cell analysis, we applied it to characterize the CpG methylation profile of HCC in mice. This HCC was initiated by the overexpression of the oncogene *c-Myc* constructed in our laboratory. Two or three tumor nodules were taken from each of the 2 HCC mice, while a corresponding part of liver was retrieved from 3 healthy mice sacrificed as the negative controls. Four technical replicates were performed with the purified genomic DNA derived from each nodule and the controls. A sum of 32 samples were processed as one healthy sample was lost during library construction. With tSNE analysis applied on either methylation levels of bins or the CpG sites, the HCC nodules and healthy livers clearly formed 3 distinct clusters (**Figure 5a**). The heat map analysis showed that the average correlation coefficients (R) were as high as 0.98 within the technical replicates of both healthy livers and HCC nodules (**Figure 5b**). In addition, mRRBS remained consistent in detecting multiple functional components in mice, and focuses on CGIs and promoters (**Supplementary Figure 6a**).

**Figure 5.**
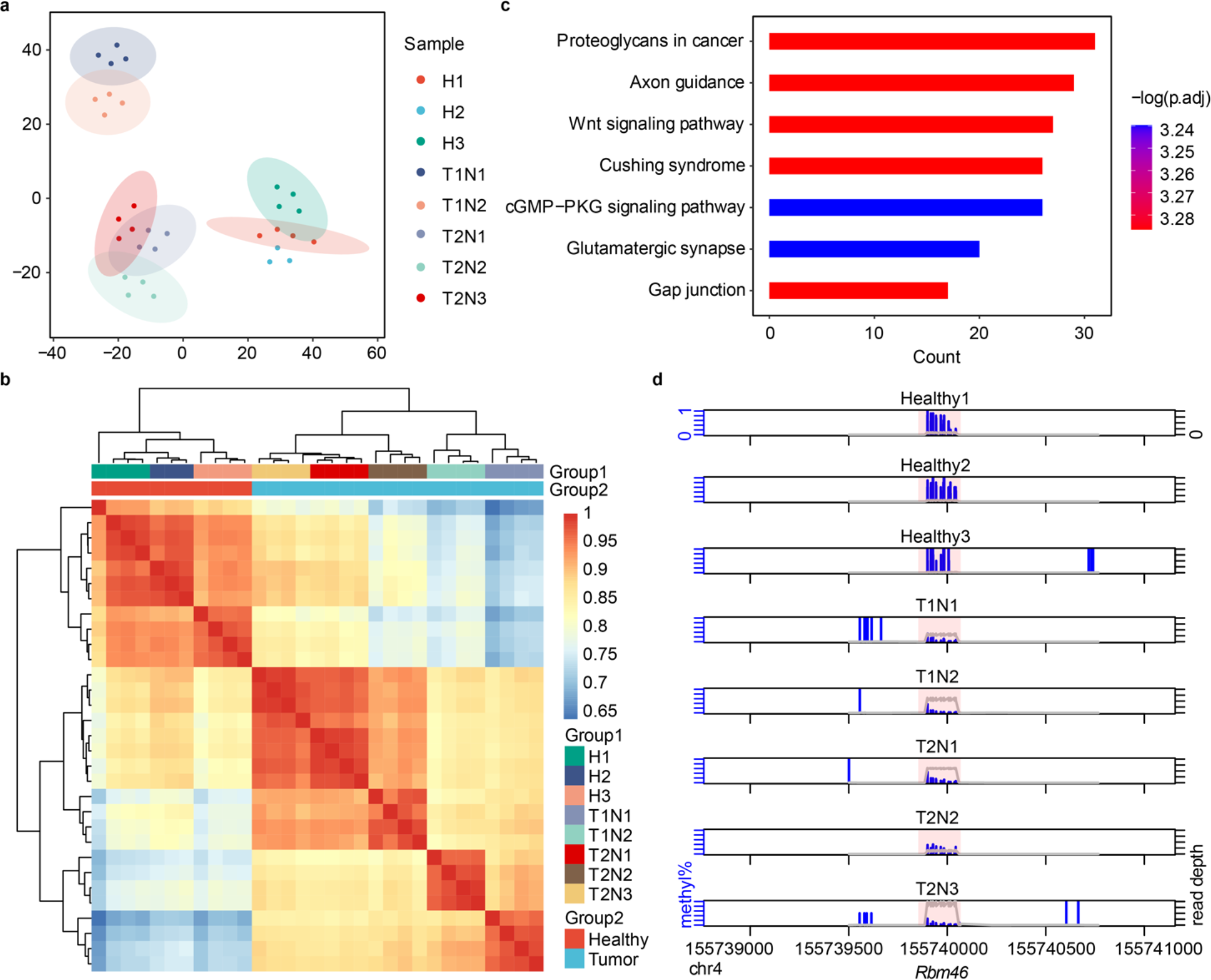
Methylation patterns of hepatocellular carcinoma (HCC) in mice uncovered by msRRBS. (a-b) Analysis of HCC bulk cells based on methylation levels in 50-kb windows over the genome for two HCC mice with two or three liver cancer nodules versus three liver biopsies from healthy mice. Except for H2 with 3 technical replicates, all the other samples were measured with 4 technical replicates. (a) Unsupervised clustering with tSNE. (b) Pearson correlation heatmap. The color key from blue to red indicates low to high correlation. (c) KEGG pathways enriched on HCC hyper-DMRs. (d) One of the HCC Hypo-DMR regions, labeled as the pink background, being highlighted by DSS package, is hypermethylated in healthy liver biopsies and hypomethylated in HCC nodules. The Y-axis from 0 to 1 represents methylation level. Blue vertical bars: methylation level. Gray line: depth of reads.

DMR analyses were performed to investigate the differential methylation status between healthy livers and HCC nodules. It was evident that the clusters were clearly identified both before (**Supplementary Figure 6b**) and after (**Supplementary Figure 6c**) the combination of technical replicates. We found that there were more DMRS in hypermethylated states than in hypomethylated states on each chromosome, and in particular, almost all DMRS in chromosome X and chromosome 3 were hypermethylated (**Supplementary Figure 6d**). KEGG analysis based on the DMRs hypermethylated in healthy livers and hypomethylated in HCC nodules highlighted signaling pathways related to the occurrence and development of liver cancer, such as axon guidance^32^ and the Wnt signaling pathway^33^ (**Figure 5c**). As examples, we highlight three DMRs on chromosomes 2, 4 and 7, where the healthy livers were hypermethylated, while all nodules of the two HCC livers were hypomethylated in *Rbm46* and *Creb5* (**Figure 5d and Supplementary Figure 6e**). Conversely, *Znf385b* displayed the opposite methylation status (**Supplementary Figure 6f**).

## DISCUSSION

DNA methylation at CpG sites regulates transcriptional activity and the stability of the genome, and is altered during differentiation, development, and aging. Single-cell analysis plays an important role in elucidating the genomic heterogeneity in multicellular organisms. In spite of other efforts to assess methylation profile free of bisulfite conversion^24, 34, 35^, single-cell bisulfite sequencing techniques have been the focus over recent years, allowing researchers to precisely analyze the epigenomic heterogeneity of single cells^3, 4, 6–10, 36^. Recent research has revealed the fundamental cellular and molecular patterns particularly in neurons and brains, embryos, stem cells, and leukemia, as well as dissecting the precise mechanism underneath DNA replication^37^. No doubt, single-cell methylation analysis methods will deepen our understanding of genomic imprinting and Fragile X Syndrome^38^, plus a variety of other fundamental processes in life, and speed up the effort for precise screening of biomarkers for cancers and other disorders. Yet, the efficiency and robustness of these techniques are still a huge challenge. In this study, we established msRRBS to analyze the methylation status of multiple single cells in-parallel at single nucleotide resolution, with inventive designs for early shortcut barcoding and pooling of the single cells. We believe that msRRBS also has the potential to work if we switch the bisulfite conversion approach to enzymatic conversion^24^.

The innovative design of msRRBS is reflected in the following aspects. Firstly, different single cells, are combined immediately after specific barcodes are added to the genome of each cell with a shortcut procedure at the beginning, so that all subsequent steps are carried out in a single tube, thus achieving multiplicity and scalability. msRRBS greatly reduces the cost, complexity and labor intensity of library construction when handling many single cells, and greatly improves the efficiency of multiple, individual cells. It is also worth noting that the whole experimental process of msRRBS library construction takes 14.5 hours, with only 4 hours of manual operation, while conventional scRRBS for a few samples takes 2-3 days and much more hands-on time.

Importantly, the data quality of msRRBS is greatly improved; this is attributed to several unique features of the method. The possible damage of the original template at the single-cell level is substantially reduced by the shortcut procedure, which is free of enzymatic DNA modifications, such as end-repair and deoxyadenosine nucleotide-adding to the MspI digested DNA fragments, in addition to the direct ligation of a short barcode-containing adapter instead of a long adapter to the MspI digested DNA fragments. This single-strand ligation avoids hard-to-remove adapter-dimers, in which the double-strand adapter is ligated to the restriction fragments by only the long strand of the adapter, while the short strand is subsequently dissociated away from the ligation product. A special non-displacement and non-nick DNA-polymerase, Sulfolobus DNA polymerase IV, is introduced to create the second strand of the long strand adapter ligated to the DNA fragments. This enzymatic single-tube process is more efficient than any physical enrichment (such as biotin-streptavidin) of the MspI digested DNA fragments, and also enables early pooling of multiple cells, taking advantage of the barcoding nucleotide combination in the adapter for each single cell. This enables the efficient amplification of the desired fragments after bisulfite conversion. These processes together minimize the possible damage of the DNA and enables the capture of the double-stranded MspI digested DNA fragments from both ends for best recovery of CpG methylation in the target fragments.

In msRRBS, the mapping rate of the raw data runs up to 79.85% with an average of 62.51% with the optimal protocol. This rate may vary with different quality of cells, associated with the genome integrity of the cells. It is similar to the latest protocols of scWGBS/scBS^10, 13, 36^, which ranges from 64-68%, and is significantly improved over the conventional scRRBS methods, which is approximately 25% ^8, 20^. Additionally in msRRBS, no cross-contamination among cells is observed, and the identity of each individual cell is traceable from the output data, ensuring the independence and reliability of the measurement for each cell. This is especially valuable for precious and rare cells, such as preimplantation embryo blastomeres, fetal cells from non-invasive prenatal testing (NIPT), and liquid biopsies of cancer, such as fine needle aspirates and circulating tumor cells.

Another advantage in the performance is that the sequencing output of msRRBS consistently enriches the key elements involved in methylation regulation, particularly CGIs and promoters. At 0.5-1 Mb reading depth, the coverage of CGIs and promoters is close to the plateau, which results in high consistency in coverage. In addition, the silico-merged data of a very limited number of single cells (i.e., pseudo-bulk built from data of 15 single cells) detects approximately equivalent CpG sites and key elements to the corresponding bulk cells. Many more DMRs are detectable from in-silico merged single cells than in real bulk cells, probably because of the independent detection of the elements in single cells, rather than the detection of the elements from the average of the bulk samples. These results favorably support the importance of a panel of single cells over a bulk of cells to detect the heterogeneity and subpopulations of a multicellular system^7, 36^.

In recent years, some efforts have been made to perform multiplex, single-cell bisulfite sequencing, mostly using the TruSeq adapter after digestion with a restriction enzyme and cell barcoding before bisulfite conversion, yet very few have succeeded. Among them, Msc-RRBS^37^ successfully employed modified TruSeq adapters with additional barcodes for multiple single-cell RRBS, and resulted in Smart-RRBS for co-analysis of RNA and CpG methylation at the single-cell level^16^. While our data was being analyzed, XRBS was reported using streptavidin to capture biotinylated adapters that are covalently ligated to (and later recovered) one strand of MspI-digested DNA fragments from the 5’ end, followed by bisulfite conversion, and the use of a semi-random primer to build the second strand for target amplification. This report detected extra elements, particularly CTCF binding motifs, beyond CGIs and promoters, with a sequencing depth much greater than RRBS^14^. Our msRRBS focuses on CGIs and promoters and uses a different strategy circumventing the labor-intensive procedures, while rescuing both strands of the MspI digested DNA fragments, minimizing DNA damage, and being flexible in fragment size recovery. Furthermore, with wider ranges of library size (switched from 175-350 bp to 175-550 bp), the coverage of CpG sites and functional elements were improved, correspondingly with a slight increase in cost of sequencing. However, after further extending the library range to 175-750 bp, the detection rate of CpG sites and elements were not significantly improved. For rare and precious cells, a 175-550 bp library range is preferred to best characterize the methylome in a single cell. Furthermore, other non-methylation sensitive restriction enzymes that enrich for CpG-rich sequence may be applied instead of MspI, and two enzymes in combination or separately for the same sample may improve the coverage^39^.

As an off-the-shelf reagent and equipment based, highly efficient, multiplexed approach, msRRBS provides a new technique to existing single-cell epigenetic technologies, primarily targeting informative CpG islands and promoter regions at genome-scale. However, msRRBS has some shortcomings and areas for further improvement. Firstly, sparsity and incomplete coverage remains an issue, due to the nature of single-cell sequencing, and because RRBS only uses MspI or its alternatives for enrichment. The unique restriction site makes the detectable CpG sites relatively limited in comparison to WGBS and XRBS. This is the downside of the advantage of high efficiency. Secondly, we have designed 96 barcodes for single-cell traceable msRRBS, but more barcodes need to be designed and validated so as to increase the throughput of msRRBS. More systematic random barcodes could be applied in combination with droplets or nanowells for truly high-throughput analysis of thousands of single cells in a run. Thirdly, the annotation of the RRBS profile of specific, single-cell subpopulations with a functional basis is a necessity and requires independent investigation.

In conclusion, we provide a highly efficient approach for medium-throughput analysis of scRRBS based on technological innovation, which is not only practically valuable when the cells are precious and rare, but also scalable to high-throughput analysis of a large number of cells in a variety of systems. This method will promote single-cell epigenomics and multiomics^9, 16, 40^, which is essential for understanding the epigenomic regulation of cellular heterogeneity.

## DATA AVAILABILITY

The public data used in this study was available in GSE149954 (XRBS), GSE27584 (bulk RRBS) and GSE86747 (bulk WGBS). Data generated from this study have been deposited to the Sequence Read Archive (SRA) under BioProject accession PRJNA945576 for unrestricted access.

## AUTHORS’ CONTRIBUTIONS

X.P. conceived and supervised the project. X.P., S.W., S.L.M., and Q.T. edited the manuscript. X.P. and L.M. designed the experiments and drafted the manuscript; L.M., Y. G., X.Y., X.L., C.C., and J. Zhou. performed the experiments; C.L. and W.P. contributed to the mouse model of HCC. Y.Z. provided the drug resistance model. Z.W., Z.L., and D.L. carried out bioinformatics analysis. X.W., J.Z., and G.L. provided variants of supports. L.M. wrote the manuscript with input from all the authors.

## CONSENT FOR PUBLICATION

All authors agree to publish this study.

---

**Figure S1.**
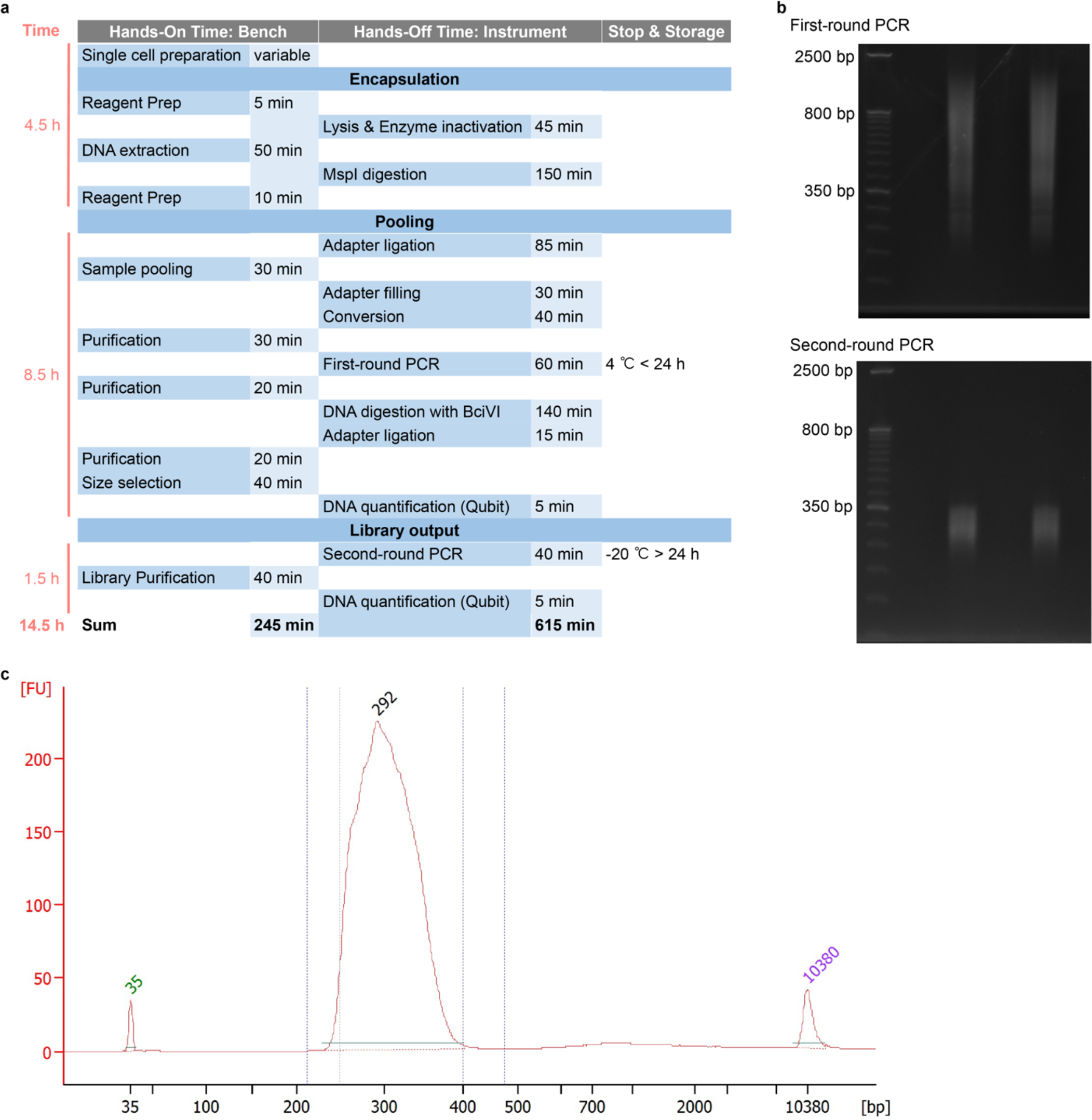
Critical technological designs with procedure time frame of msRRBS. (a) The standard duration of each step of msRRBS procedure. (b) E-Gel™ EX 2% agarose gel electropherograms of the first and second rounds of PCR amplification. (c) Bioanalyzer electropherograms of msRRBS libraries. Here the library fragments are >175 bp with a mean length near 300 bp.

**Figure S2.**
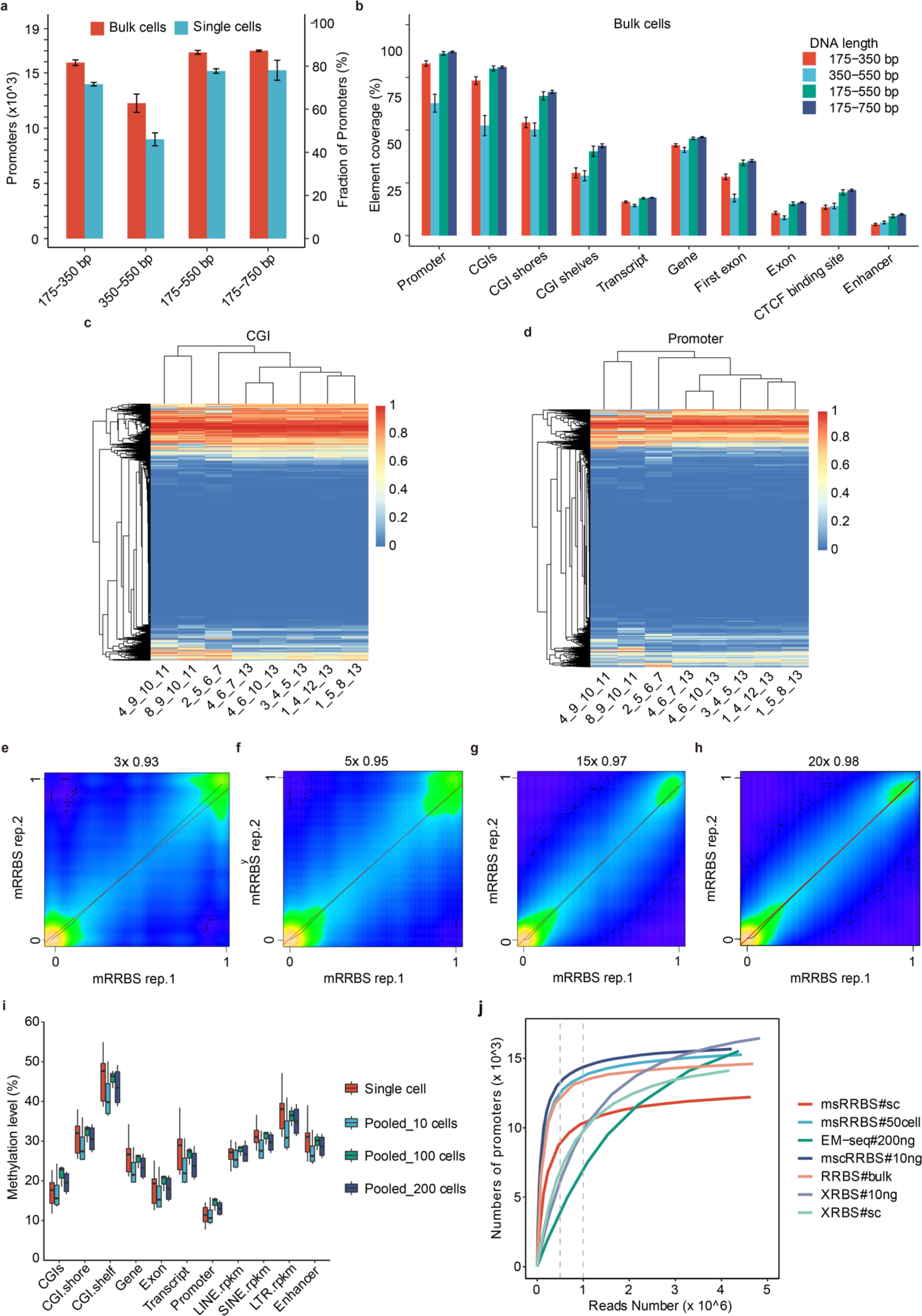
Mapping rates and element coverages of msRRBS with different ranges of library size assayed in cell line K562. (a-b) In the single-cell sample, except for the library size of 175-750bp, which is 4 samples, the remaining three libraries contained only 3 samples. (a) Detection rate of promoters associated with different library sizes in single cells and bulk cells. (b) Coverage of various functional elements along the genome associated with different library sizes, detected in K562 bulk cells (n=4). (c-d) Heatmap shows the methylation levels of CGIs (a) and promoters (b) in four randomly in silico-merged single cells from cell line K562. The number on the x axis represents the barcode of the merged single cells. (e-h) Pearson correlation heatmap compares the methylation pattern based on CpG sites, acquired by mRRBS, between technical replicates of bulk cells for cell line K562 in coverage ≥ 3× (c, R = 0.93), ≥ 5× (d, R = 0.95), ≥ 15× (e, R = 0.97) and ≥ 20× (f, R = 0.98). (i) Methylation levels in different genomic regions of single cells (n=15), micro-bulk cells of 10, 100, and 200 cells (n=2) of K562. (j) Number of promoters containing CpG islands with at least 1 CpG site covered as a function of sequencing depth for msRRBS, mRRBS, traditional RRBS, EM-Seq and XRBS in cell line K562. The vertical gray dashed lines denote 0.5 M and 1 M clean reads as a threshold for comparison.

**Figure S3.**
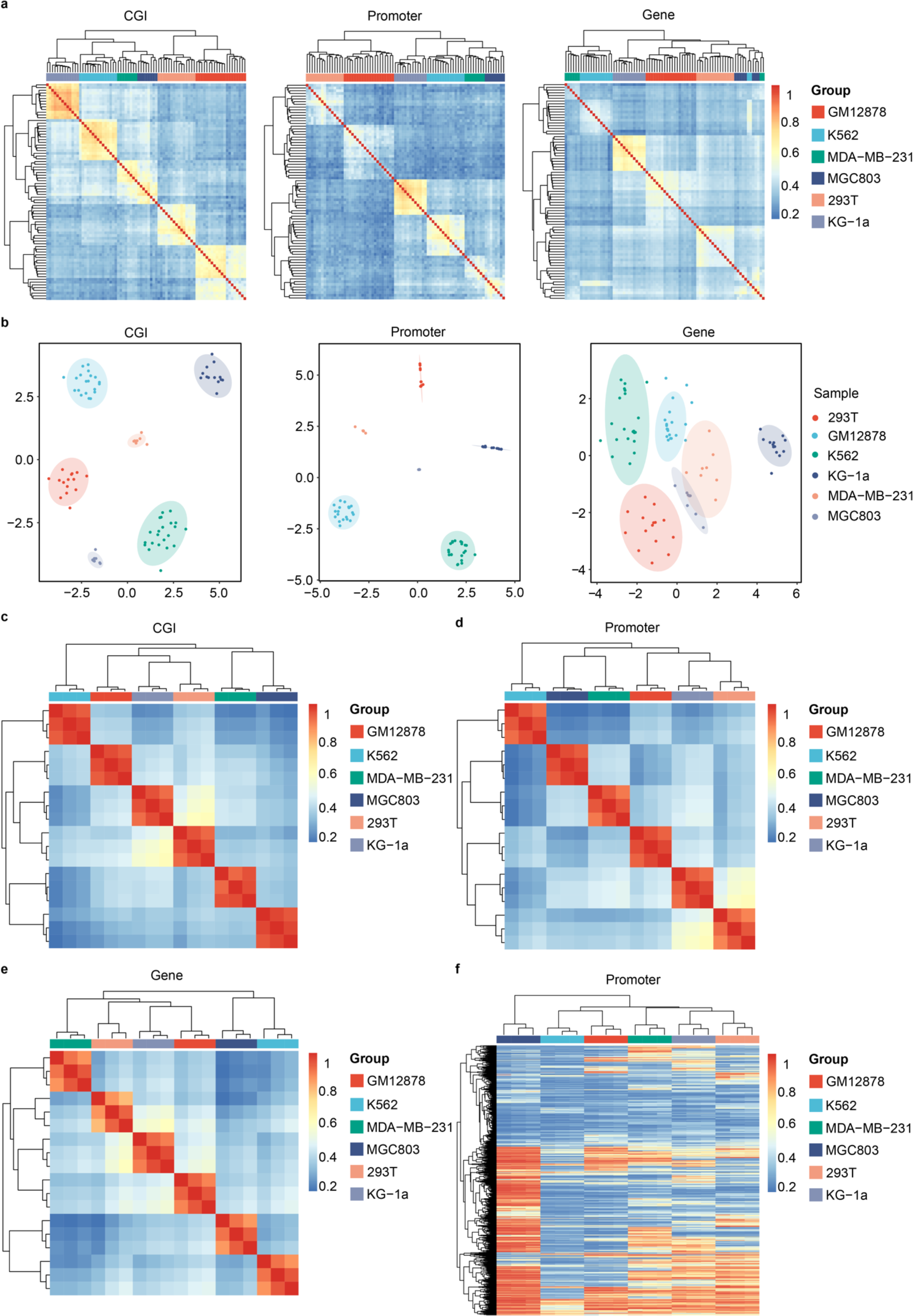
Distinct methylation profiling of six cell lines in variant functional elements examined by msRRBS. (a) Pearson correlation heatmap shows the methylation levels of CGIs, promoters, and genes among the single-cell samples. The color key from blue to red indicates low to high correlation. Six cell lines: 293T (n=15), GM12878 (n=20), K562 (n=15), KG-1a (n=13), MDA-MB-231 (n=8) and MGC803 (n=8) were analyzed. (b) Unsupervised clustering of the single cells from the six cell lines based on methylation levels of CGIs, promoters, and genes, generated with tSNE. (c-e) Pearson correlation heatmap shows the methylation levels of CGIs (c), promoters (d), and genes (e) of 2, 4, and 8 single cells in the corresponding region were randomly in silico-merged from each of the six cell lines. (f) Methylation profiles of 2, 4, and 8 single cells in the promoter region were randomly in silico-merged for each of the six cell lines. The number 0 represents unmethylated, and the number 1 represents fully methylated. The color changes from blue to red to indicate a gradual increase in methylation level.

**Figure S4.**
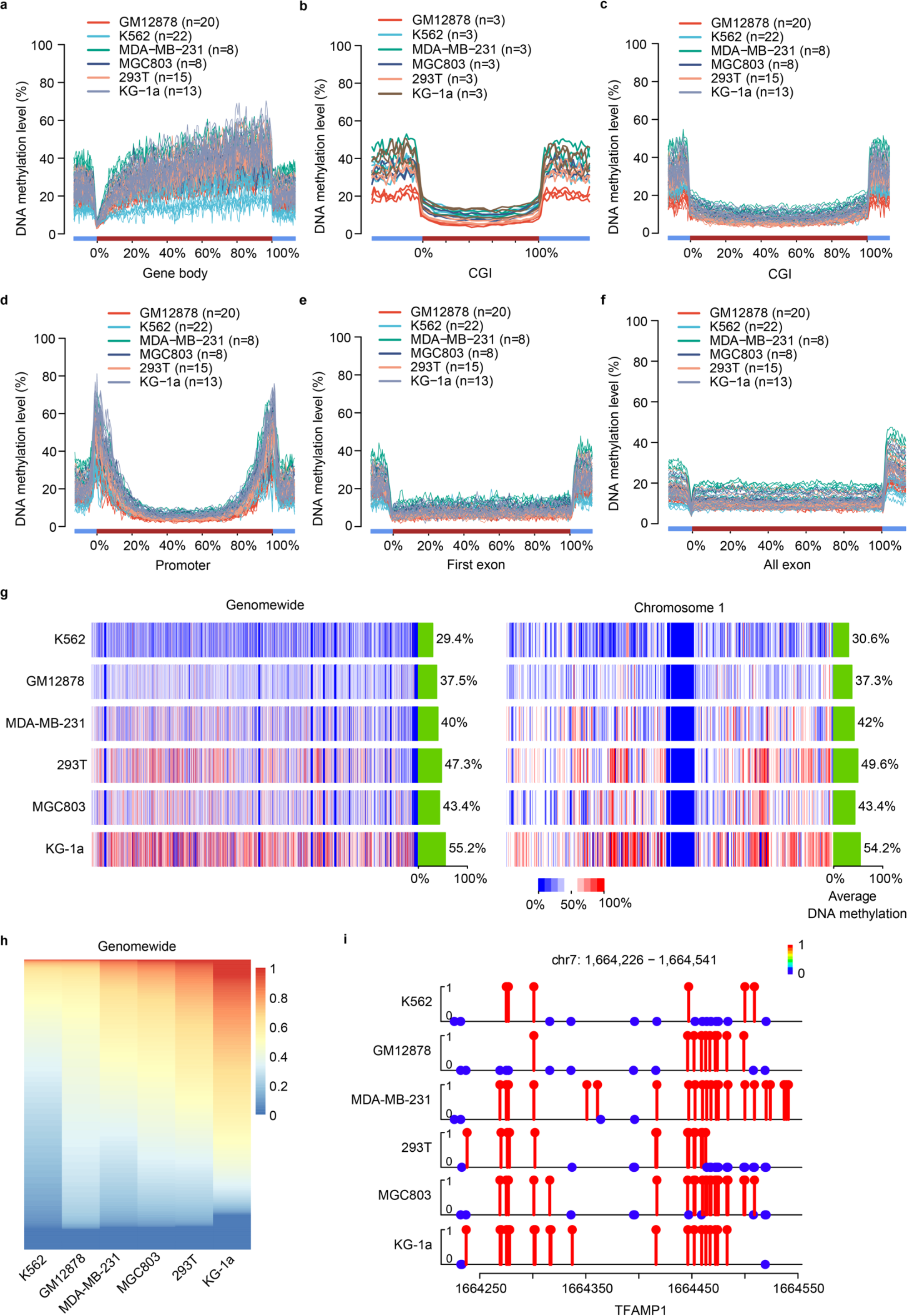
Reliability and reproducibility of msRRBS in the measurement of methylation pattern. (a-f) DNAme patterns across gene body (a), CGIs (b and c), promoters (d), first exons (e), all exons (f) and 5 kb upstream and downstream of the TSSs and the TESs of all RefSeq genes in combination. (g-h) DNAme levels of whole genome (g and h, the methylation levels of figure h are sorted) and chromosome 1 (g, right) for each cell line with 8 single cells in silico-merged by windowing method with at least 20 CpG sites per 50-kb bin. (i) Methylation status of six cell lines in a representative window with gene TFAMP1 (an example arbitrarily selected). The methylation levels of most of the CpG sites of KG-1a in the window are fully methylated (Red lollipop), and most of the CpG sites of K562 in the window are unmethylated (blue circles).

**Figure S5.**
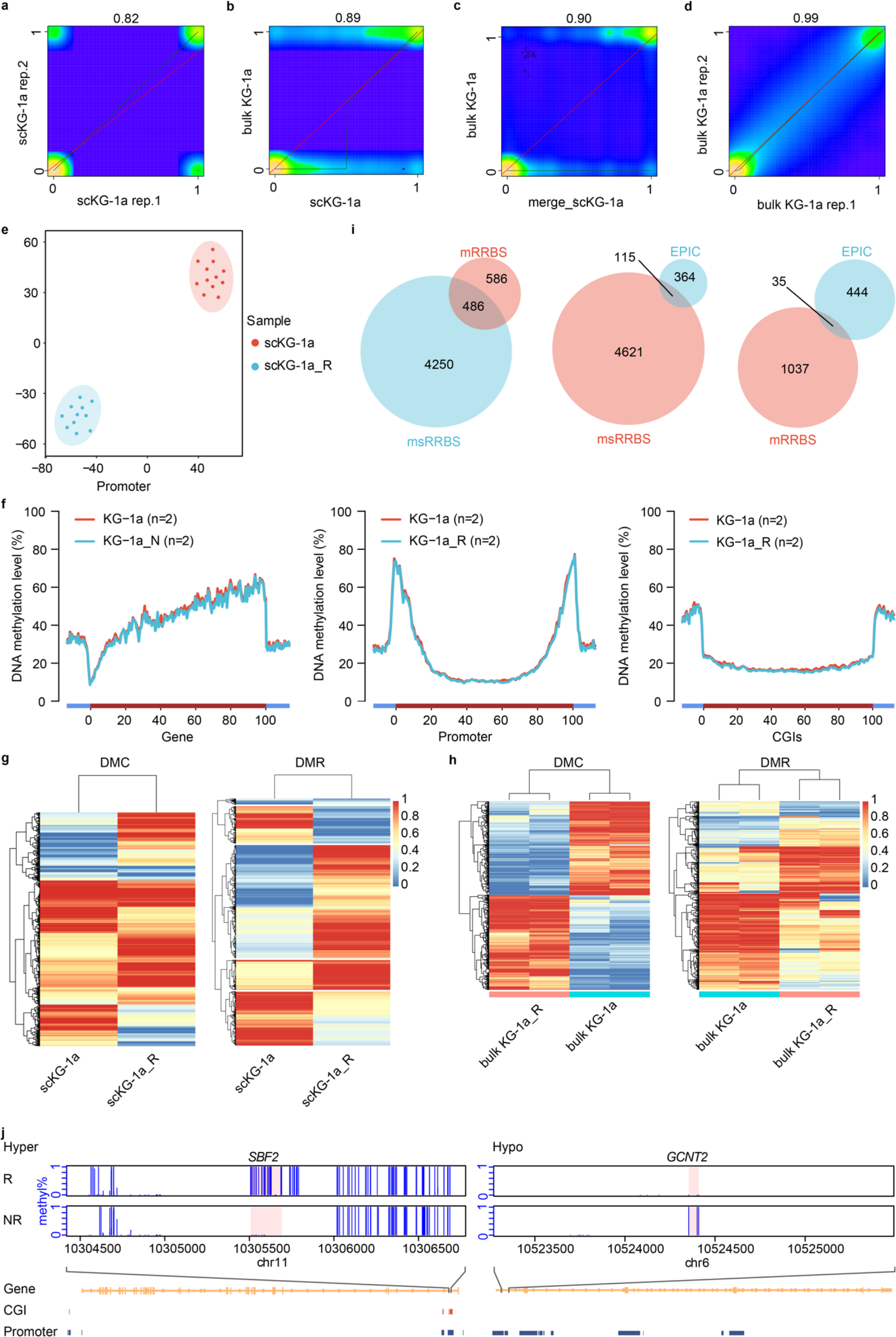
msRRBS study on KG-1a resistant to cytarabine (KG-1a_R) versus the original KG-1a. (a-d) Pearson correlation heatmap compares methylation values of individual CpG sites acquired by msRRBS between two single cells (a, R = 0.82), between a single cell and a bulk of cells (b, R = 0.89), between pseudo-bulk (merged 13 cells) and a bulk of cells (c, 0.90), and between technical replicates of bulk cells (d, R = 0.99) for the original culture of cell line KG-1a. (e) Unsupervised clustering of single cells based on the methylation of promoters for KG-1a_R (n=11) and KG-1a (n=12), generated by tSNE. (f) Average DNAme patterns across gene bodies, promoters, and CGIs from the data set of bulk cells by mRRBS. (g-j) KG-1a_R versus KG-1a cells. (g) DMC methylation heatmap for in silico-merged single cells of KG-1a. (h) DMC and DMR methylation heatmap for bulk cells from KG-1a_R versus KG-1a. (i) Hypo-DMR intersections of KG-1a_R vs. KG-1a among methods msRRBS, mRRBS and EPIC methylation array. (j) Two windows each contain a DMR of merged single cells from KG-1a_R versus the corresponding KG-1a population (MWU-test p-value < 0.05, number of CpG sites in one DMR ≥ 3, the distance between two independent DMRs ≥ 100bp, length of DMR ≥ 50bp).

**Figure S6.**
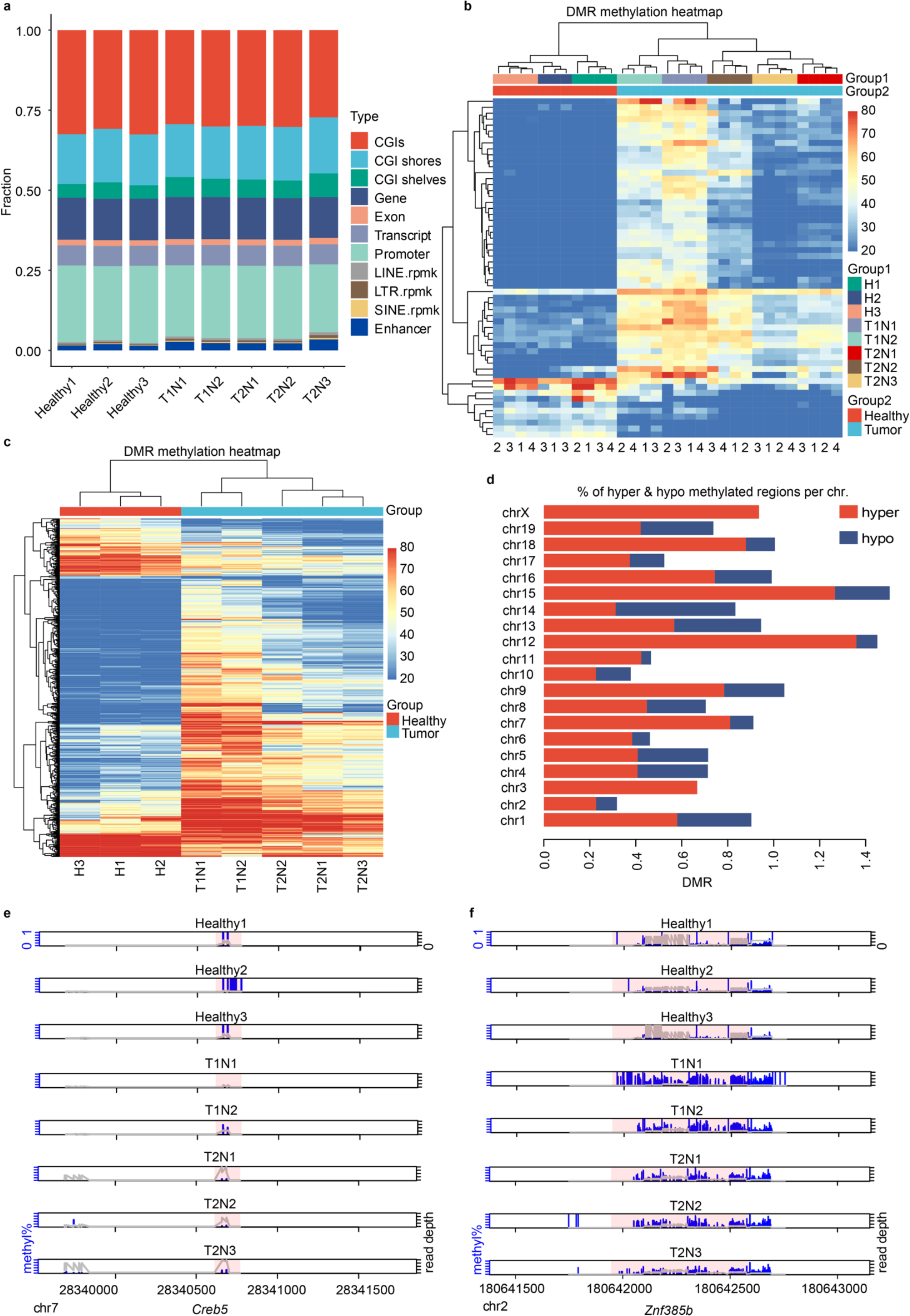
Additional features and DMR examples of mRRBS analysis for the hepatocellular carcinoma (HCC) in mice. (a) Average detection rates of different genomic regions in each sample. (b-c) DMR methylation heatmap for individual samples (b) or in silico-merge of technical replicates (c). Each sample was measured with 3 or 4 technical replicates. (e-f) Two representative DMRs between HCC and liver from healthy mice (MWU-test p-value < 0.05, number of CpG sites in one DMR ≥ 3, the distance between two independent DMRs ≥ 100bp, length of DMR ≥ 50bp). (d) Percentage of hyper & hypo methylated regions per chromosome, q value<0.01 & methylation diff. >=25 %. (e) *Creb5* is hypermethylated in healthy mice and hypomethylated in HCC mice. (f) *Znf385b* is hypomethylated in healthy mice and hypermethylated in HCC mice. The number 0 represents unmethylated, and the number 1 represents methylated. The pink background represents the length of the DMR region. The blue vertical bars represent methylation levels, and higher vertical bars represent higher methylation levels. The higher the gray line is, the deeper the reads are.

## References

1. Lister, R. et al. Human DNA methylomes at base resolution show widespread epigenomic differences. Nature 462, 315–322 (2009).

2. Meissner, A. et al. Reduced representation bisulfite sequencing for comparative high-resolution DNA methylation analysis. Nucleic Acids Res 33, 5868–5877 (2005).

3. Farlik, M. et al. DNA Methylation Dynamics of Human Hematopoietic Stem Cell Differentiation. Cell Stem Cell 19, 808–822 (2016).

4. Luo, C. et al. Single-cell methylomes identify neuronal subtypes and regulatory elements in mammalian cortex. Science 357, 600–604 (2017).

5. Nichols, R.V. et al. High-throughput robust single-cell DNA methylation profiling with sciMETv2. Nat Commun 13, 7627 (2022).

6. Gaiti, F. et al. Epigenetic evolution and lineage histories of chronic lymphocytic leukaemia. Nature 569, 576–580 (2019).

7. Liu, H. et al. DNA methylation atlas of the mouse brain at single-cell resolution. Nature 598, 120–128 (2021).

8. Guo, H. et al. Single-cell methylome landscapes of mouse embryonic stem cells and early embryos analyzed using reduced representation bisulfite sequencing. Genome Res 23, 2126–2135 (2013).

9. Angermueller, C. et al. Parallel single-cell sequencing links transcriptional and epigenetic heterogeneity. Nat Methods 13, 229–232 (2016).

10. Smallwood, S.A. et al. Single-cell genome-wide bisulfite sequencing for assessing epigenetic heterogeneity. Nat Methods 11, 817–820 (2014).

11. Miura, F., Enomoto, Y., Dairiki, R. & Ito, T. Amplification-free whole-genome bisulfite sequencing by post-bisulfite adaptor tagging. Nucleic Acids Res 40, e136 (2012).

12. Farlik, M. et al. Single-cell DNA methylome sequencing and bioinformatic inference of epigenomic cell-state dynamics. Cell Rep 10, 1386–1397 (2015).

13. Luo, C. et al. Robust single-cell DNA methylome profiling with snmC-seq2. Nat Commun 9, 3824 (2018).

14. Shareef, S.J. et al. Extended-representation bisulfite sequencing of gene regulatory elements in multiplexed samples and single cells. Nat Biotechnol 39, 1086–1094 (2021).

15. Smith, Z.D. et al. Epigenetic restriction of extraembryonic lineages mirrors the somatic transition to cancer. Nature 549, 543–547 (2017).

16. Gu, H. et al. Smart-RRBS for single-cell methylome and transcriptome analysis. Nat Protoc 16, 4004–4030 (2021).

17. Martin, M. Cutadapt removes adapter sequences from high-throughput sequencing reads. Embnet Journal 17 (2011).

18. Bolger, A.M., Lohse, M. & Usadel, B. Trimmomatic: a flexible trimmer for Illumina sequence data. Bioinformatics 30, 2114–2120 (2014).

19. Krueger, F. & Andrews, S.R. Bismark: a flexible aligner and methylation caller for Bisulfite-Seq applications. Bioinformatics 27, 1571–1572 (2011).

20. Guo, H. et al. Profiling DNA methylome landscapes of mammalian cells with single-cell reduced-representation bisulfite sequencing. Nat Protoc 10, 645–659 (2015).

21. Guo, W. et al. CGmapTools improves the precision of heterozygous SNV calls and supports allele-specific methylation detection and visualization in bisulfite-sequencing data. Bioinformatics 34, 381–387 (2018).

22. Wu, H. et al. Detection of differentially methylated regions from whole-genome bisulfite sequencing data without replicates. Nucleic Acids Res 43, e141 (2015).

23. Heinz, S. et al. Simple combinations of lineage-determining transcription factors prime cis-regulatory elements required for macrophage and B cell identities. Mol Cell 38, 576–589 (2010).

24. Vaisvila, R. et al. Enzymatic methyl sequencing detects DNA methylation at single-base resolution from picograms of DNA. Genome Res 31, 1280–1289 (2021).

25. Milella, M. et al. Synergistic induction of apoptosis by simultaneous disruption of the Bcl-2 and MEK/MAPK pathways in acute myelogenous leukemia. Blood 99, 3461–3464 (2002).

26. Shao, M. et al. Inhibition of Calcium Signaling Prevents Exhaustion and Enhances Anti-Leukemia Efficacy of CAR-T Cells via SOCE-Calcineurin-NFAT and Glycolysis Pathways. Adv Sci (Weinh) 9, e2103508 (2022).

27. Dunwell, T.L. et al. Frequent epigenetic inactivation of the SLIT2 gene in chronic and acute lymphocytic leukemia. Epigenetics 4, 265–269 (2009).

28. Zhang, H. et al. Engagement of I-branching {beta}-1, 6-N-acetylglucosaminyltransferase 2 in breast cancer metastasis and TGF-{beta} signaling. Cancer Res 71, 4846–4856 (2011).

29. Nicosia, L. et al. Pharmacological inhibition of LSD1 triggers myeloid differentiation by targeting GSE1 oncogenic functions in AML. Oncogene 41, 878–894 (2022).

30. Yamakawa, K. et al. DSCAM: a novel member of the immunoglobulin superfamily maps in a Down syndrome region and is involved in the development of the nervous system. Hum Mol Genet 7, 227–237 (1998).

31. Lv, J. et al. High expression of long non-coding RNA SBF2-AS1 promotes proliferation in non-small cell lung cancer. J Exp Clin Cancer Res 35, 75 (2016).

32. Liang, J.Q. et al. Dietary cholesterol promotes steatohepatitis related hepatocellular carcinoma through dysregulated metabolism and calcium signaling. Nat Commun 9, 4490 (2018).

33. Huang, G. et al. CircRNA hsa_circRNA_104348 promotes hepatocellular carcinoma progression through modulating miR-187-3p/RTKN2 axis and activating Wnt/β-catenin pathway. Cell Death Dis 11, 1065 (2020).

34. Han, L. et al. Bisulfite-independent analysis of CpG island methylation enables genome-scale stratification of single cells. Nucleic Acids Res 45, e77 (2017).

35. Viswanathan, R., Cheruba, E. & Cheow, L.F. DNA Analysis by Restriction Enzyme (DARE) enables concurrent genomic and epigenomic characterization of single cells. Nucleic Acids Res 47, e122 (2019).

36. Mulqueen, R.M. et al. Highly scalable generation of DNA methylation profiles in single cells. Nat Biotechnol 36, 428–431 (2018).

37. Charlton, J. et al. Global delay in nascent strand DNA methylation. Nat Struct Mol Biol 25, 327–332 (2018).

38. Yasukochi, Y. et al. X chromosome-wide analyses of genomic DNA methylation states and gene expression in male and female neutrophils. Proc Natl Acad Sci U S A 107, 3704–3709 (2010).

39. Wang, J. et al. Double restriction-enzyme digestion improves the coverage and accuracy of genome-wide CpG methylation profiling by reduced representation bisulfite sequencing. BMC Genomics 14, 11 (2013).

40. Li, L. et al. Single-cell multi-omics sequencing of human early embryos. Nat Cell Biol 20, 847–858 (2018).

