## Supplementary Table for "Shortcut barcoding and early pooling for scalable multiplex single-cell reduced-representation CpG methylation sequencing at single nucleotide resolution"

### Supplementary Table 1. List of 96 adapter with barcodes (#1-28 adapters are validated).

| msRRBS Adapter | Long strand | Long strand in IDT ordering format (5' -> 3') | Inline Barcode | Short strand | Short strand in IDT ordering format (5' -> 3') |
|---|---|---|---|---|---|
| 1 | AdBcL1 | AAG TAG GTA T/iMe-dC/iMe-dC/ GTG AGT GGT G [6 bases barcode] | AAGAAT | AdBcS1 | CG [6 bases barcode (Reverse complementation)] CACCA/3Amino/ |
| 2 | AdBcL2 | AAG TAG GTA T/iMe-dC/iMe-dC/ GTG AGT GGT G [6 bases barcode] | TAAGTA | AdBcS2 | CG [6 bases barcode (Reverse complementation)] CACCA/3Amino/ |
| 3 | AdBcL3 | AAG TAG GTA T/iMe-dC/iMe-dC/ GTG AGT GGT G [6 bases barcode] | AGTTAT | AdBcS3 | CG [6 bases barcode (Reverse complementation)] CACCA/3Amino/ |
| 4 | AdBcL4 | AAG TAG GTA T/iMe-dC/iMe-dC/ GTG AGT GGT G [6 bases barcode] | TAATGT | AdBcS4 | CG [6 bases barcode (Reverse complementation)] CACCA/3Amino/ |
| 5 | AdBcL5 | AAG TAG GTA T/iMe-dC/iMe-dC/ GTG AGT GGT G [6 bases barcode] | TTAGAA | AdBcS5 | CG [6 bases barcode (Reverse complementation)] CACCA/3Amino/ |
| 6 | AdBcL6 | AAG TAG GTA T/iMe-dC/iMe-dC/ GTG AGT GGT G [6 bases barcode] | ATGTAA | AdBcS6 | CG [6 bases barcode (Reverse complementation)] CACCA/3Amino/ |
| 7 | AdBcL7 | AAG TAG GTA T/iMe-dC/iMe-dC/ GTG AGT GGT G [6 bases barcode] | ATGTAA | AdBcS7 | CG [6 bases barcode (Reverse complementation)] CACCA/3Amino/ |
| 8 | AdBcL8 | AAG TAG GTA T/iMe-dC/iMe-dC/ GTG AGT GGT G [6 bases barcode] | ATAGTT | AdBcS8 | CG [6 bases barcode (Reverse complementation)] CACCA/3Amino/ |
| 9 | AdBcL9 | AAG TAG GTA T/iMe-dC/iMe-dC/ GTG AGT GGT G [6 bases barcode] | AAGTTA | AdBcS9 | CG [6 bases barcode (Reverse complementation)] CACCA/3Amino/ |
| 10 | AdBcL10 | AAG TAG GTA T/iMe-dC/iMe-dC/ GTG AGT GGT G [6 bases barcode] | TATGAT | AdBcS10 | CG [6 bases barcode (Reverse complementation)] CACCA/3Amino/ |
| 11 | AdBcL11 | AAG TAG GTA T/iMe-dC/iMe-dC/ GTG AGT GGT G [6 bases barcode] | AGAATA | AdBcS11 | CG [6 bases barcode (Reverse complementation)] CACCA/3Amino/ |
| 12 | AdBcL12 | AAG TAG GTA T/iMe-dC/iMe-dC/ GTG AGT GGT G [6 bases barcode] | TATAGA | AdBcS12 | CG [6 bases barcode (Reverse complementation)] CACCA/3Amino/ |
| 13 | AdBcL13 | AAG TAG GTA T/iMe-dC/iMe-dC/ GTG AGT GGT G [6 bases barcode] | TTGATT | AdBcS13 | CG [6 bases barcode (Reverse complementation)] CACCA/3Amino/ |
| 14 | AdBcL14 | AAG TAG GTA T/iMe-dC/iMe-dC/ GTG AGT GGT G [6 bases barcode] | TTTGAT | AdBcS14 | CG [6 bases barcode (Reverse complementation)] CACCA/3Amino/ |
| 15 | AdBcL15 | AAG TAG GTA T/iMe-dC/iMe-dC/ GTG AGT GGT G [6 bases barcode] | TGATAT | AdBcS15 | CG [6 bases barcode (Reverse complementation)] CACCA/3Amino/ |
| 16 | AdBcL16 | AAG TAG GTA T/iMe-dC/iMe-dC/ GTG AGT GGT G [6 bases barcode] | TATGAA | AdBcS16 | CG [6 bases barcode (Reverse complementation)] CACCA/3Amino/ |
| 17 | AdBcL17 | AAG TAG GTA T/iMe-dC/iMe-dC/ GTG AGT GGT G [6 bases barcode] | AGATAG | AdBcS17 | CG [6 bases barcode (Reverse complementation)] CACCA/3Amino/ |
| 18 | AdBcL18 | AAG TAG GTA T/iMe-dC/iMe-dC/ GTG AGT GGT G [6 bases barcode] | AGGATG | AdBcS18 | CG [6 bases barcode (Reverse complementation)] CACCA/3Amino/ |
| 19 | AdBcL19 | AAG TAG GTA T/iMe-dC/iMe-dC/ GTG AGT GGT G [6 bases barcode] | AGGTTA | AdBcS19 | CG [6 bases barcode (Reverse complementation)] CACCA/3Amino/ |
| 20 | AdBcL20 | AAG TAG GTA T/iMe-dC/iMe-dC/ GTG AGT GGT G [6 bases barcode] | AGGTAA | AdBcS20 | CG [6 bases barcode (Reverse complementation)] CACCA/3Amino/ |
| 21 | AdBcL21 | AAG TAG GTA T/iMe-dC/iMe-dC/ GTG AGT GGT G [6 bases barcode] | AGGGAT | AdBcS21 | CG [6 bases barcode (Reverse complementation)] CACCA/3Amino/ |
| 22 | AdBcL22 | AAG TAG GTA T/iMe-dC/iMe-dC/ GTG AGT GGT G [6 bases barcode] | GAGAAT | AdBcS22 | CG [6 bases barcode (Reverse complementation)] CACCA/3Amino/ |
| 23 | AdBcL23 | AAG TAG GTA T/iMe-dC/iMe-dC/ GTG AGT GGT G [6 bases barcode] | TGTGAG | AdBcS23 | CG [6 bases barcode (Reverse complementation)] CACCA/3Amino/ |
| 24 | AdBcL24 | AAG TAG GTA T/iMe-dC/iMe-dC/ GTG AGT GGT G [6 bases barcode] | TTAAAG | AdBcS24 | CG [6 bases barcode (Reverse complementation)] CACCA/3Amino/ |
| 25 | AdBcL25 | AAG TAG GTA T/iMe-dC/iMe-dC/ GTG AGT GGT G [6 bases barcode] | GAATTG | AdBcS25 | CG [6 bases barcode (Reverse complementation)] CACCA/3Amino/ |
| 26 | AdBcL26 | AAG TAG GTA T/iMe-dC/iMe-dC/ GTG AGT GGT G [6 bases barcode] | AGGAGT | AdBcS26 | CG [6 bases barcode (Reverse complementation)] CACCA/3Amino/ |
| 27 | AdBcL27 | AAG TAG GTA T/iMe-dC/iMe-dC/ GTG AGT GGT G [6 bases barcode] | GAGATA | AdBcS27 | CG [6 bases barcode (Reverse complementation)] CACCA/3Amino/ |
| 28 | AdBcL28 | AAG TAG GTA T/iMe-dC/iMe-dC/ GTG AGT GGT G [6 bases barcode] | GAGGTA | AdBcS28 | CG [6 bases barcode (Reverse complementation)] CACCA/3Amino/ |
| 29 | AdBcL29 | AAG TAG GTA T/iMe-dC/iMe-dC/ GTG AGT GGT G [6 bases barcode] | AGTGAG | AdBcS29 | CG [6 bases barcode (Reverse complementation)] CACCA/3Amino/ |
| 30 | AdBcL30 | AAG TAG GTA T/iMe-dC/iMe-dC/ GTG AGT GGT G [6 bases barcode] | ATTGAG | AdBcS30 | CG [6 bases barcode (Reverse complementation)] CACCA/3Amino/ |
| 31 | AdBcL31 | AAG TAG GTA T/iMe-dC/iMe-dC/ GTG AGT GGT G [6 bases barcode] | AGAGTG | AdBcS31 | CG [6 bases barcode (Reverse complementation)] CACCA/3Amino/ |
| 32 | AdBcL32 | AAG TAG GTA T/iMe-dC/iMe-dC/ GTG AGT GGT G [6 bases barcode] | TTGAGG | AdBcS32 | CG [6 bases barcode (Reverse complementation)] CACCA/3Amino/ |
| 33 | AdBcL33 | AAG TAG GTA T/iMe-dC/iMe-dC/ GTG AGT GGT G [6 bases barcode] | AGGATT | AdBcS33 | CG [6 bases barcode (Reverse complementation)] CACCA/3Amino/ |
| 34 | AdBcL34 | AAG TAG GTA T/iMe-dC/iMe-dC/ GTG AGT GGT G [6 bases barcode] | TAGTAT | AdBcS34 | CG [6 bases barcode (Reverse complementation)] CACCA/3Amino/ |
| 35 | AdBcL35 | AAG TAG GTA T/iMe-dC/iMe-dC/ GTG AGT GGT G [6 bases barcode] | TGTAAT | AdBcS35 | CG [6 bases barcode (Reverse complementation)] CACCA/3Amino/ |
| 36 | AdBcL36 | AAG TAG GTA T/iMe-dC/iMe-dC/ GTG AGT GGT G [6 bases barcode] | TGAGTT | AdBcS36 | CG [6 bases barcode (Reverse complementation)] CACCA/3Amino/ |
| 37 | AdBcL37 | AAG TAG GTA T/iMe-dC/iMe-dC/ GTG AGT GGT G [6 bases barcode] | TGGAGA | AdBcS37 | CG [6 bases barcode (Reverse complementation)] CACCA/3Amino/ |
| 38 | AdBcL38 | AAG TAG GTA T/iMe-dC/iMe-dC/ GTG AGT GGT G [6 bases barcode] | TGATTG | AdBcS38 | CG [6 bases barcode (Reverse complementation)] CACCA/3Amino/ |
| 39 | AdBcL39 | AAG TAG GTA T/iMe-dC/iMe-dC/ GTG AGT GGT G [6 bases barcode] | TGTTGA | AdBcS39 | CG [6 bases barcode (Reverse complementation)] CACCA/3Amino/ |
| 40 | AdBcL40 | AAG TAG GTA T/iMe-dC/iMe-dC/ GTG AGT GGT G [6 bases barcode] | GTATGA | AdBcS40 | CG [6 bases barcode (Reverse complementation)] CACCA/3Amino/ |

| | | | | | |
|---|---|---|---|---|---|
| 41 | AdBcL41 | AAG TAG GTA T/iMe-dC/iMe-dC/ GTG AGT GGT G [6 bases barcode] | TGGATG | AdBcS41 | CG [6 bases barcode (Reverse complementation)] CACCA/3Amino/ |
| 42 | AdBcL42 | AAG TAG GTA T/iMe-dC/iMe-dC/ GTG AGT GGT G [6 bases barcode] | TTGTGA | AdBcS42 | CG [6 bases barcode (Reverse complementation)] CACCA/3Amino/ |
| 43 | AdBcL43 | AAG TAG GTA T/iMe-dC/iMe-dC/ GTG AGT GGT G [6 bases barcode] | GTAGTA | AdBcS43 | CG [6 bases barcode (Reverse complementation)] CACCA/3Amino/ |
| 44 | AdBcL44 | AAG TAG GTA T/iMe-dC/iMe-dC/ GTG AGT GGT G [6 bases barcode] | AAGTGT | AdBcS44 | CG [6 bases barcode (Reverse complementation)] CACCA/3Amino/ |
| 45 | AdBcL45 | AAG TAG GTA T/iMe-dC/iMe-dC/ GTG AGT GGT G [6 bases barcode] | GGTTAA | AdBcS45 | CG [6 bases barcode (Reverse complementation)] CACCA/3Amino/ |
| 46 | AdBcL46 | AAG TAG GTA T/iMe-dC/iMe-dC/ GTG AGT GGT G [6 bases barcode] | AGTATG | AdBcS46 | CG [6 bases barcode (Reverse complementation)] CACCA/3Amino/ |
| 47 | AdBcL47 | AAG TAG GTA T/iMe-dC/iMe-dC/ GTG AGT GGT G [6 bases barcode] | AGAGAT | AdBcS47 | CG [6 bases barcode (Reverse complementation)] CACCA/3Amino/ |
| 48 | AdBcL48 | AAG TAG GTA T/iMe-dC/iMe-dC/ GTG AGT GGT G [6 bases barcode] | ATAGAG | AdBcS48 | CG [6 bases barcode (Reverse complementation)] CACCA/3Amino/ |
| 49 | AdBcL49 | AAG TAG GTA T/iMe-dC/iMe-dC/ GTG AGT GGT G [6 bases barcode] | ATAAGA | AdBcS49 | CG [6 bases barcode (Reverse complementation)] CACCA/3Amino/ |
| 50 | AdBcL50 | AAG TAG GTA T/iMe-dC/iMe-dC/ GTG AGT GGT G [6 bases barcode] | ATAATG | AdBcS50 | CG [6 bases barcode (Reverse complementation)] CACCA/3Amino/ |
| 51 | AdBcL51 | AAG TAG GTA T/iMe-dC/iMe-dC/ GTG AGT GGT G [6 bases barcode] | TGTAGG | AdBcS51 | CG [6 bases barcode (Reverse complementation)] CACCA/3Amino/ |
| 52 | AdBcL52 | AAG TAG GTA T/iMe-dC/iMe-dC/ GTG AGT GGT G [6 bases barcode] | TAGAAG | AdBcS52 | CG [6 bases barcode (Reverse complementation)] CACCA/3Amino/ |
| 53 | AdBcL53 | AAG TAG GTA T/iMe-dC/iMe-dC/ GTG AGT GGT G [6 bases barcode] | TGAGAG | AdBcS53 | CG [6 bases barcode (Reverse complementation)] CACCA/3Amino/ |
| 54 | AdBcL54 | AAG TAG GTA T/iMe-dC/iMe-dC/ GTG AGT GGT G [6 bases barcode] | TAGAGT | AdBcS54 | CG [6 bases barcode (Reverse complementation)] CACCA/3Amino/ |
| 55 | AdBcL55 | AAG TAG GTA T/iMe-dC/iMe-dC/ GTG AGT GGT G [6 bases barcode] | GAGTAA | AdBcS55 | CG [6 bases barcode (Reverse complementation)] CACCA/3Amino/ |
| 56 | AdBcL56 | AAG TAG GTA T/iMe-dC/iMe-dC/ GTG AGT GGT G [6 bases barcode] | TAATAG | AdBcS56 | CG [6 bases barcode (Reverse complementation)] CACCA/3Amino/ |
| 57 | AdBcL57 | AAG TAG GTA T/iMe-dC/iMe-dC/ GTG AGT GGT G [6 bases barcode] | TATATG | AdBcS57 | CG [6 bases barcode (Reverse complementation)] CACCA/3Amino/ |
| 58 | AdBcL58 | AAG TAG GTA T/iMe-dC/iMe-dC/ GTG AGT GGT G [6 bases barcode] | AGGGTA | AdBcS58 | CG [6 bases barcode (Reverse complementation)] CACCA/3Amino/ |
| 59 | AdBcL59 | AAG TAG GTA T/iMe-dC/iMe-dC/ GTG AGT GGT G [6 bases barcode] | GAGTGG | AdBcS59 | CG [6 bases barcode (Reverse complementation)] CACCA/3Amino/ |
| 60 | AdBcL60 | AAG TAG GTA T/iMe-dC/iMe-dC/ GTG AGT GGT G [6 bases barcode] | GAGGAT | AdBcS60 | CG [6 bases barcode (Reverse complementation)] CACCA/3Amino/ |
| 61 | AdBcL61 | AAG TAG GTA T/iMe-dC/iMe-dC/ GTG AGT GGT G [6 bases barcode] | TGTGGA | AdBcS61 | CG [6 bases barcode (Reverse complementation)] CACCA/3Amino/ |
| 62 | AdBcL62 | AAG TAG GTA T/iMe-dC/iMe-dC/ GTG AGT GGT G [6 bases barcode] | TGTATA | AdBcS62 | CG [6 bases barcode (Reverse complementation)] CACCA/3Amino/ |
| 63 | AdBcL63 | AAG TAG GTA T/iMe-dC/iMe-dC/ GTG AGT GGT G [6 bases barcode] | TGTTAG | AdBcS63 | CG [6 bases barcode (Reverse complementation)] CACCA/3Amino/ |
| 64 | AdBcL64 | AAG TAG GTA T/iMe-dC/iMe-dC/ GTG AGT GGT G [6 bases barcode] | TGAATG | AdBcS64 | CG [6 bases barcode (Reverse complementation)] CACCA/3Amino/ |
| 65 | AdBcL65 | AAG TAG GTA T/iMe-dC/iMe-dC/ GTG AGT GGT G [6 bases barcode] | TGAAGT | AdBcS65 | CG [6 bases barcode (Reverse complementation)] CACCA/3Amino/ |
| 66 | AdBcL66 | AAG TAG GTA T/iMe-dC/iMe-dC/ GTG AGT GGT G [6 bases barcode] | TGGTAT | AdBcS66 | CG [6 bases barcode (Reverse complementation)] CACCA/3Amino/ |
| 67 | AdBcL67 | AAG TAG GTA T/iMe-dC/iMe-dC/ GTG AGT GGT G [6 bases barcode] | TGGTTA | AdBcS67 | CG [6 bases barcode (Reverse complementation)] CACCA/3Amino/ |
| 68 | AdBcL68 | AAG TAG GTA T/iMe-dC/iMe-dC/ GTG AGT GGT G [6 bases barcode] | TGGAAT | AdBcS68 | CG [6 bases barcode (Reverse complementation)] CACCA/3Amino/ |
| 69 | AdBcL69 | AAG TAG GTA T/iMe-dC/iMe-dC/ GTG AGT GGT G [6 bases barcode] | TGGATA | AdBcS69 | CG [6 bases barcode (Reverse complementation)] CACCA/3Amino/ |
| 70 | AdBcL70 | AAG TAG GTA T/iMe-dC/iMe-dC/ GTG AGT GGT G [6 bases barcode] | AGGAAT | AdBcS70 | CG [6 bases barcode (Reverse complementation)] CACCA/3Amino/ |
| 71 | AdBcL71 | AAG TAG GTA T/iMe-dC/iMe-dC/ GTG AGT GGT G [6 bases barcode] | TAGTTG | AdBcS71 | CG [6 bases barcode (Reverse complementation)] CACCA/3Amino/ |
| 72 | AdBcL72 | AAG TAG GTA T/iMe-dC/iMe-dC/ GTG AGT GGT G [6 bases barcode] | TAGGTT | AdBcS72 | CG [6 bases barcode (Reverse complementation)] CACCA/3Amino/ |
| 73 | AdBcL73 | AAG TAG GTA T/iMe-dC/iMe-dC/ GTG AGT GGT G [6 bases barcode] | TAGGAA | AdBcS73 | CG [6 bases barcode (Reverse complementation)] CACCA/3Amino/ |
| 74 | AdBcL74 | AAG TAG GTA T/iMe-dC/iMe-dC/ GTG AGT GGT G [6 bases barcode] | TGAGTG | AdBcS74 | CG [6 bases barcode (Reverse complementation)] CACCA/3Amino/ |
| 75 | AdBcL75 | AAG TAG GTA T/iMe-dC/iMe-dC/ GTG AGT GGT G [6 bases barcode] | AATGAA | AdBcS75 | CG [6 bases barcode (Reverse complementation)] CACCA/3Amino/ |
| 76 | AdBcL76 | AAG TAG GTA T/iMe-dC/iMe-dC/ GTG AGT GGT G [6 bases barcode] | AAATGG | AdBcS76 | CG [6 bases barcode (Reverse complementation)] CACCA/3Amino/ |
| 77 | AdBcL77 | AAG TAG GTA T/iMe-dC/iMe-dC/ GTG AGT GGT G [6 bases barcode] | AAGTTG | AdBcS77 | CG [6 bases barcode (Reverse complementation)] CACCA/3Amino/ |
| 78 | AdBcL78 | AAG TAG GTA T/iMe-dC/iMe-dC/ GTG AGT GGT G [6 bases barcode] | GTATAG | AdBcS78 | CG [6 bases barcode (Reverse complementation)] CACCA/3Amino/ |
| 79 | AdBcL79 | AAG TAG GTA T/iMe-dC/iMe-dC/ GTG AGT GGT G [6 bases barcode] | ATTATA | AdBcS79 | CG [6 bases barcode (Reverse complementation)] CACCA/3Amino/ |
| 80 | AdBcL80 | AAG TAG GTA T/iMe-dC/iMe-dC/ GTG AGT GGT G [6 bases barcode] | ATTAAT | AdBcS80 | CG [6 bases barcode (Reverse complementation)] CACCA/3Amino/ |
| 81 | AdBcL81 | AAG TAG GTA T/iMe-dC/iMe-dC/ GTG AGT GGT G [6 bases barcode] | ATTGTA | AdBcS81 | CG [6 bases barcode (Reverse complementation)] CACCA/3Amino/ |
| 82 | AdBcL82 | AAG TAG GTA T/iMe-dC/iMe-dC/ GTG AGT GGT G [6 bases barcode] | ATTTAG | AdBcS82 | CG [6 bases barcode (Reverse complementation)] CACCA/3Amino/ |
| 83 | AdBcL83 | AAG TAG GTA T/iMe-dC/iMe-dC/ GTG AGT GGT G [6 bases barcode] | ATTTGA | AdBcS83 | CG [6 bases barcode (Reverse complementation)] CACCA/3Amino/ |

| | | | | | |
|---|---|---|---|---|---|
| 84 | AdBcL84 | AAG TAG GTA T/iMe-dC/iMe-dC/ GTG AGT GGT G [6 bases barcode] | TTTGTA | AdBcS84 | CG [6 bases barcode (Reverse complementation)] CACCA/3Amino/ |
| 85 | AdBcL85 | AAG TAG GTA T/iMe-dC/iMe-dC/ GTG AGT GGT G [6 bases barcode] | TTTAGG | AdBcS85 | CG [6 bases barcode (Reverse complementation)] CACCA/3Amino/ |
| 86 | AdBcL86 | AAG TAG GTA T/iMe-dC/iMe-dC/ GTG AGT GGT G [6 bases barcode] | TTTGAG | AdBcS86 | CG [6 bases barcode (Reverse complementation)] CACCA/3Amino/ |
| 87 | AdBcL87 | AAG TAG GTA T/iMe-dC/iMe-dC/ GTG AGT GGT G [6 bases barcode] | TTTGGA | AdBcS87 | CG [6 bases barcode (Reverse complementation)] CACCA/3Amino/ |
| 88 | AdBcL88 | AAG TAG GTA T/iMe-dC/iMe-dC/ GTG AGT GGT G [6 bases barcode] | GAATGA | AdBcS88 | CG [6 bases barcode (Reverse complementation)] CACCA/3Amino/ |
| 89 | AdBcL89 | AAG TAG GTA T/iMe-dC/iMe-dC/ GTG AGT GGT G [6 bases barcode] | GAATAG | AdBcS89 | CG [6 bases barcode (Reverse complementation)] CACCA/3Amino/ |
| 90 | AdBcL90 | AAG TAG GTA T/iMe-dC/iMe-dC/ GTG AGT GGT G [6 bases barcode] | GAAGAT | AdBcS90 | CG [6 bases barcode (Reverse complementation)] CACCA/3Amino/ |
| 91 | AdBcL91 | AAG TAG GTA T/iMe-dC/iMe-dC/ GTG AGT GGT G [6 bases barcode] | GAAGTA | AdBcS91 | CG [6 bases barcode (Reverse complementation)] CACCA/3Amino/ |
| 92 | AdBcL92 | AAG TAG GTA T/iMe-dC/iMe-dC/ GTG AGT GGT G [6 bases barcode] | TTAGGA | AdBcS92 | CG [6 bases barcode (Reverse complementation)] CACCA/3Amino/ |
| 93 | AdBcL93 | AAG TAG GTA T/iMe-dC/iMe-dC/ GTG AGT GGT G [6 bases barcode] | TTAGAG | AdBcS93 | CG [6 bases barcode (Reverse complementation)] CACCA/3Amino/ |
| 94 | AdBcL94 | AAG TAG GTA T/iMe-dC/iMe-dC/ GTG AGT GGT G [6 bases barcode] | ATAGGA | AdBcS94 | CG [6 bases barcode (Reverse complementation)] CACCA/3Amino/ |
| 95 | AdBcL95 | AAG TAG GTA T/iMe-dC/iMe-dC/ GTG AGT GGT G [6 bases barcode] | ATATGG | AdBcS95 | CG [6 bases barcode (Reverse complementation)] CACCA/3Amino/ |
| 96 | AdBcL96 | AAG TAG GTA T/iMe-dC/iMe-dC/ GTG AGT GGT G [6 bases barcode] | TATAAT | AdBcS96 | CG [6 bases barcode (Reverse complementation)] CACCA/3Amino/ |

Note: Adapter #1-28 were validated, preferably #1-16, and more preferably #7, #9, #12, #15.

Supplementary Table 2. The primers used in the study.

| Name | sequence (5′–3′) | Index |
|---|---|---|
| PreAP | AAGTAGGTATCCGTGAGTGGTG | N/A |
| AdSp1 | 5Phos/GATCGGAAGAGCACACGTCTGAACTCCAGTCAC | N/A |
| AdSp2 | ACACTCTTTCCCTACACGACGCTCTTCCGATC*T | N/A |
| P5 | AATGATACGGCGACCACCGAGATCTACACTCTTTCCCTACACGACG*C | N/A |
| P7-1 | CAAGCAGAAGACGGCATACGAGAT[6 bases]GTGACTGGAGTTCAGACGTGTGCTCTTCCGATC*T | ATCACG |
| P7-2 | CAAGCAGAAGACGGCATACGAGAT[6 bases]GTGACTGGAGTTCAGACGTGTGCTCTTCCGATC*T | CGATGT |
| P7-3 | CAAGCAGAAGACGGCATACGAGAT[6 bases]GTGACTGGAGTTCAGACGTGTGCTCTTCCGATC*T | TTAGGC |
| P7-4 | CAAGCAGAAGACGGCATACGAGAT[6 bases]GTGACTGGAGTTCAGACGTGTGCTCTTCCGATC*T | TGACCA |
| P7-5 | CAAGCAGAAGACGGCATACGAGAT[6 bases]GTGACTGGAGTTCAGACGTGTGCTCTTCCGATC*T | ACAGTG |
| P7-6 | CAAGCAGAAGACGGCATACGAGAT[6 bases]GTGACTGGAGTTCAGACGTGTGCTCTTCCGATC*T | GCCAAT |
| P7-7 | CAAGCAGAAGACGGCATACGAGAT[6 bases]GTGACTGGAGTTCAGACGTGTGCTCTTCCGATC*T | CAGATC |
| P7-8 | CAAGCAGAAGACGGCATACGAGAT[6 bases]GTGACTGGAGTTCAGACGTGTGCTCTTCCGATC*T | ACTTGA |
| P7-9 | CAAGCAGAAGACGGCATACGAGAT[6 bases]GTGACTGGAGTTCAGACGTGTGCTCTTCCGATC*T | GATCAG |
| P7-10 | CAAGCAGAAGACGGCATACGAGAT[6 bases]GTGACTGGAGTTCAGACGTGTGCTCTTCCGATC*T | TAGCTT |
| P7-11 | CAAGCAGAAGACGGCATACGAGAT[6 bases]GTGACTGGAGTTCAGACGTGTGCTCTTCCGATC*T | GGCTAC |
| P7-12 | CAAGCAGAAGACGGCATACGAGAT[6 bases]GTGACTGGAGTTCAGACGTGTGCTCTTCCGATC*T | CTTGTA |
| P7-13 | CAAGCAGAAGACGGCATACGAGAT[6 bases]GTGACTGGAGTTCAGACGTGTGCTCTTCCGATC*T | AGTCAA |
| P7-14 | CAAGCAGAAGACGGCATACGAGAT[6 bases]GTGACTGGAGTTCAGACGTGTGCTCTTCCGATC*T | AGTTCC |
| P7-15 | CAAGCAGAAGACGGCATACGAGAT[6 bases]GTGACTGGAGTTCAGACGTGTGCTCTTCCGATC*T | ATGTCA |
| P7-16 | CAAGCAGAAGACGGCATACGAGAT[6 bases]GTGACTGGAGTTCAGACGTGTGCTCTTCCGATC*T | CCGTCC |
| P7-17 | CAAGCAGAAGACGGCATACGAGAT[6 bases]GTGACTGGAGTTCAGACGTGTGCTCTTCCGATC*T | GTAGAG |
| P7-18 | CAAGCAGAAGACGGCATACGAGAT[6 bases]GTGACTGGAGTTCAGACGTGTGCTCTTCCGATC*T | GTCCGC |
| P7-19 | CAAGCAGAAGACGGCATACGAGAT[6 bases]GTGACTGGAGTTCAGACGTGTGCTCTTCCGATC*T | GTGAAA |
| P7-20 | CAAGCAGAAGACGGCATACGAGAT[6 bases]GTGACTGGAGTTCAGACGTGTGCTCTTCCGATC*T | GTGGCC |
| P7-21 | CAAGCAGAAGACGGCATACGAGAT[6 bases]GTGACTGGAGTTCAGACGTGTGCTCTTCCGATC*T | GTTTCG |
| P7-22 | CAAGCAGAAGACGGCATACGAGAT[6 bases]GTGACTGGAGTTCAGACGTGTGCTCTTCCGATC*T | CGTACG |
| P7-23 | CAAGCAGAAGACGGCATACGAGAT[6 bases]GTGACTGGAGTTCAGACGTGTGCTCTTCCGATC*T | GAGTGG |
| P7-24 | CAAGCAGAAGACGGCATACGAGAT[6 bases]GTGACTGGAGTTCAGACGTGTGCTCTTCCGATC*T | GGTAGC |
| P7-25 | CAAGCAGAAGACGGCATACGAGAT[6 bases]GTGACTGGAGTTCAGACGTGTGCTCTTCCGATC*T | ACTGAT |
| P7-26 | CAAGCAGAAGACGGCATACGAGAT[6 bases]GTGACTGGAGTTCAGACGTGTGCTCTTCCGATC*T | ATGAGC |
| P7-27 | CAAGCAGAAGACGGCATACGAGAT[6 bases]GTGACTGGAGTTCAGACGTGTGCTCTTCCGATC*T | ATTCCT |
| P7-28 | CAAGCAGAAGACGGCATACGAGAT[6 bases]GTGACTGGAGTTCAGACGTGTGCTCTTCCGATC*T | CAAAAG |
| P7-29 | CAAGCAGAAGACGGCATACGAGAT[6 bases]GTGACTGGAGTTCAGACGTGTGCTCTTCCGATC*T | CAACTA |
| P7-30 | CAAGCAGAAGACGGCATACGAGAT[6 bases]GTGACTGGAGTTCAGACGTGTGCTCTTCCGATC*T | CACCGG |
| P7-31 | CAAGCAGAAGACGGCATACGAGAT[6 bases]GTGACTGGAGTTCAGACGTGTGCTCTTCCGATC*T | CACGAT |
| P7-32 | CAAGCAGAAGACGGCATACGAGAT[6 bases]GTGACTGGAGTTCAGACGTGTGCTCTTCCGATC*T | CACTCA |
| P7-33 | CAAGCAGAAGACGGCATACGAGAT[6 bases]GTGACTGGAGTTCAGACGTGTGCTCTTCCGATC*T | CAGGCG |
| P7-34 | CAAGCAGAAGACGGCATACGAGAT[6 bases]GTGACTGGAGTTCAGACGTGTGCTCTTCCGATC*T | CATGGC |
| P7-35 | CAAGCAGAAGACGGCATACGAGAT[6 bases]GTGACTGGAGTTCAGACGTGTGCTCTTCCGATC*T | CATTTT |
| P7-36 | CAAGCAGAAGACGGCATACGAGAT[6 bases]GTGACTGGAGTTCAGACGTGTGCTCTTCCGATC*T | CCAACA |
| P7-37 | CAAGCAGAAGACGGCATACGAGAT[6 bases]GTGACTGGAGTTCAGACGTGTGCTCTTCCGATC*T | CGGAAT |

| | | |
|---|---|---|
| P7-38 | CAAGCAGAAGACGGCATACGAGAT[6 bases]GTGACTGGAGTTCAGACGTGTGCTCTTCCGATC*T | CTAGCT |
| P7-39 | CAAGCAGAAGACGGCATACGAGAT[6 bases]GTGACTGGAGTTCAGACGTGTGCTCTTCCGATC*T | CTATAC |
| P7-40 | CAAGCAGAAGACGGCATACGAGAT[6 bases]GTGACTGGAGTTCAGACGTGTGCTCTTCCGATC*T | CTCAGA |
| P7-41 | CAAGCAGAAGACGGCATACGAGAT[6 bases]GTGACTGGAGTTCAGACGTGTGCTCTTCCGATC*T | GACGAC |
| P7-42 | CAAGCAGAAGACGGCATACGAGAT[6 bases]GTGACTGGAGTTCAGACGTGTGCTCTTCCGATC*T | TAATCG |
| P7-43 | CAAGCAGAAGACGGCATACGAGAT[6 bases]GTGACTGGAGTTCAGACGTGTGCTCTTCCGATC*T | TACAGC |
| P7-44 | CAAGCAGAAGACGGCATACGAGAT[6 bases]GTGACTGGAGTTCAGACGTGTGCTCTTCCGATC*T | TATAAT |
| P7-45 | CAAGCAGAAGACGGCATACGAGAT[6 bases]GTGACTGGAGTTCAGACGTGTGCTCTTCCGATC*T | TCATTC |
| P7-46 | CAAGCAGAAGACGGCATACGAGAT[6 bases]GTGACTGGAGTTCAGACGTGTGCTCTTCCGATC*T | TCCCGA |
| P7-47 | CAAGCAGAAGACGGCATACGAGAT[6 bases]GTGACTGGAGTTCAGACGTGTGCTCTTCCGATC*T | TCGAAG |
| P7-48 | CAAGCAGAAGACGGCATACGAGAT[6 bases]GTGACTGGAGTTCAGACGTGTGCTCTTCCGATC*T | TCGGCA |

Note: * Phosphorothioate linkage

Supplementary Table 3. Summary of the unique CpG sites covered at 1×, 3×, 5× and 10× in bulk of K562 (representative samples).

| Samples | Unique CpGs (1×) | Unique CpGs (3×) | Unique CpGs (5×) | Unique CpGs (10×) | CGIs (1×) | Conversion rate (%) | Alignment rate (%) |
|---|---|---|---|---|---|---|---|
| Pooled_10 cells | 3,563,769 | 2,822,256 | 2,516,921 | 2,081,295 | 22,293 | 99.66 | 70.00 |
| Pooled_ 50 cells | 3,154,710 | 2,172,666 | 1,738,712 | 1,176,008 | 21,716 | 99.72 | 77.51 |
| Pooled_100 cells | 3,013,602 | 2,407,262 | 2,151,063 | 1,777,244 | 21,528 | 99.59 | 73.26 |
| Pooled_200 cells | 3,917,791 | 3,102,392 | 2,078,247 | 2,078,247 | 22,970 | 99.64 | 82.13 |
| Pooled_10 ng DNA | 4,058,568 | 2,771,449 | 2,257,520 | 1,613,248 | 23,474 | 99.57 | 78.30 |
| Pooled_50 ng DNA | 4,115,442 | 2,848,831 | 2,343,323 | 1,696,442 | 23,697 | 99.55 | 78.33 |
| Pooled_100 ng DNA | 4,093,477 | 2,776,011 | 2,193,685 | 1,449,033 | 23,685 | 99.91 | 77.07 |
| Pooled_200 ng DNA | 4,038,380 | 2,776,476 | 2,249,235 | 1,585,663 | 23,863 | 99.86 | 74.88 |

Supplementary Table 4. Summary of conversion rate, mapping rate and the unique covered CpG sites and elements in single cells and bulks of K562 with different library size.

| Samples | Conversion rate (%) | Mapping rate (%) | CpGs | %met CHG | %met CHH | Promoters | Promoter (%) | CGIs | CGIs (%) | CGI shores | CGI shores (%) | CGI shelves | CGI shelves (%) | Transcript | Transcript (%) | Gene | Gene (%) | First exon | First exon (%) | Exon | Exon (%) | CTCF binding site | CTCF binding site (%) | Enhancer | Enhancer (%) |
|---|---|---|---|---|---|---|---|---|---|---|---|---|---|---|---|---|---|---|---|---|---|---|---|---|---|
| 175-350bp#sc1 | 99.59 | 60.02 | 1,897,734 | 0.41 | 0.41 | 13,810 | 70.77% | 18,604 | 66.44% | 22,807 | 40.49% | 9,379 | 19.83% | 31,977 | 13.87% | 23,614 | 38.94% | 12,463 | 20.57% | 46,459 | 7.20% | 16,137 | 9.17% | 4,406 | 3.44% |
| 175-350bp#sc2 | 99.57 | 53.12 | 1,870,601 | 0.43 | 0.43 | 13,989 | 71.69% | 18,571 | 66.33% | 22,675 | 40.26% | 9,123 | 19.29% | 31,949 | 13.85% | 23,620 | 38.95% | 12,693 | 20.95% | 46,200 | 7.16% | 15,906 | 9.04% | 4,303 | 3.36% |
| 175-350bp#sc3 | 99.59 | 58.63 | 2,010,679 | 0.40 | 0.41 | 14,126 | 72.39% | 18,835 | 67.27% | 23,209 | 41.21% | 9,360 | 19.79% | 32,435 | 14.06% | 23,958 | 39.51% | 12,870 | 21.24% | 46,961 | 7.28% | 16,451 | 9.35% | 4,590 | 3.59% |
| 350-550bp#sc1 | 99.57 | 58.16 | 1,364,526 | 0.41 | 0.40 | 9,137 | 46.83% | 10,963 | 39.15% | 20,799 | 36.93% | 9,471 | 20.02% | 27,600 | 11.97% | 21,538 | 35.52% | 6,956 | 11.48% | 34,442 | 5.34% | 16,230 | 9.23% | 5,094 | 3.98% |
| 350-550bp#sc2 | 99.59 | 59.31 | 1,079,644 | 0.41 | 0.42 | 8,342 | 42.75% | 9,656 | 34.49% | 18,427 | 32.72% | 8,177 | 17.29% | 25,686 | 11.14% | 20,371 | 33.60% | 6,088 | 10.05% | 28,944 | 4.49% | 13,745 | 7.81% | 4,366 | 3.41% |
| 350-550bp#sc3 | 99.60 | 59.00 | 1,453,211 | 0.41 | 0.40 | 9,477 | 48.57% | 11,451 | 40.90% | 21,526 | 38.22% | 9,684 | 20.47% | 28,124 | 12.20% | 21,886 | 36.09% | 7,368 | 12.16% | 35,684 | 5.53% | 16,818 | 9.56% | 5,341 | 4.17% |
| 175-550bp#sc1 (merged) | 99.59 | 58.68 | 2,979,962 | 0.41 | 0.41 | 15,127 | 77.52% | 20,813 | 74.33% | 30,267 | 53.74% | 13,859 | 29.30% | 36,029 | 15.62% | 25,788 | 42.53% | 15,594 | 25.73% | 66,392 | 10.29% | 25,281 | 14.37% | 7,764 | 6.07% |
| 175-550bp#sc2 (merged) | 99.58 | 56.74 | 2,677,400 | 0.42 | 0.42 | 14,984 | 76.79% | 20,255 | 72.34% | 28,896 | 51.30% | 12,799 | 27.06% | 35,297 | 15.31% | 25,391 | 41.88% | 15,158 | 25.01% | 62,059 | 9.62% | 23,265 | 13.23% | 7,088 | 5.54% |
| 175-550bp#sc3 (merged) | 99.60 | 58.89 | 3,156,994 | 0.41 | 0.40 | 15,389 | 78.87% | 21,131 | 75.47% | 30,916 | 54.89% | 13,989 | 29.57% | 36,463 | 15.81% | 26,076 | 43.00% | 16,164 | 26.68% | 67,707 | 10.49% | 25,909 | 14.73% | 8,050 | 6.29% |
| 175-750bp#sc1 | 99.61 | 57.42 | 1,991,431 | 0.41 | 0.41 | 13,919 | 71.33% | 18,551 | 66.25% | 25,558 | 45.38% | 11,195 | 23.67% | 33,136 | 14.37% | 24,316 | 40.10% | 12,916 | 21.31% | 52,419 | 8.12% | 19,485 | 11.08% | 5,725 | 4.47% |
| 175-750bp#sc2 | 99.59 | 65.59 | 3,478,493 | 0.41 | 0.41 | 15,709 | 80.51% | 21,711 | 77.54% | 31,589 | 56.09% | 14,596 | 30.85% | 37,215 | 16.14% | 26,335 | 43.43% | 16,850 | 27.81% | 71,162 | 11.03% | 26,469 | 15.05% | 8,045 | 6.29% |
| 175-750bp#sc3 | 99.60 | 61.63 | 3,116,922 | 0.43 | 0.43 | 15,389 | 78.87% | 21,209 | 75.75% | 30,107 | 53.46% | 13,737 | 29.04% | 36,572 | 15.86% | 26,044 | 42.95% | 16,132 | 26.62% | 66,366 | 10.28% | 24,680 | 14.03% | 7,673 | 6.00% |
| 175-750bp#sc4 | 99.60 | 65.37 | 3,791,693 | 0.41 | 0.41 | 15,929 | 81.63% | 22,383 | 79.94% | 32,725 | 58.10% | 15,469 | 32.70% | 37,861 | 16.42% | 26,690 | 44.02% | 17,486 | 28.86% | 74,904 | 11.61% | 28,083 | 15.97% | 8,759 | 6.85% |
| 175-350bp#bulk1 | 99.69 | 77.29 | 3,648,021 | 0.35 | 0.33 | 15,850 | 81.23% | 22,801 | 81.43% | 30,143 | 53.52% | 14,088 | 29.78% | 36,794 | 15.95% | 25,929 | 42.76% | 16,594 | 27.38% | 68,661 | 10.64% | 23,527 | 13.38% | 6,766 | 5.29% |
| 175-350bp#bulk2 | 99.69 | 77.89 | 3,332,468 | 0.34 | 0.33 | 15,690 | 80.41% | 22,303 | 79.65% | 28,909 | 51.33% | 13,150 | 27.80% | 36,199 | 15.70% | 25,631 | 42.27% | 16,166 | 26.68% | 64,680 | 10.02% | 22,137 | 12.59% | 6,201 | 4.85% |
| 175-350bp#bulk3 | 99.64 | 78.64 | 4,546,088 | 0.36 | 0.35 | 16,282 | 83.44% | 23,647 | 84.45% | 32,135 | 57.06% | 15,660 | 33.10% | 38,379 | 16.64% | 26,615 | 43.89% | 17,992 | 29.69% | 76,020 | 11.78% | 26,028 | 14.80% | 7,740 | 6.05% |
| 175-350bp#bulk4 | 99.72 | 77.12 | 3,710,974 | 0.34 | 0.33 | 15,878 | 81.37% | 22,843 | 81.58% | 29,702 | 52.74% | 13,453 | 28.44% | 36,747 | 15.93% | 25,860 | 42.65% | 16,810 | 27.74% | 68,102 | 10.55% | 23,015 | 13.09% | 6,507 | 5.09% |
| 350-550bp#bulk1 | 99.54 | 60.02 | 2,679,255 | 0.34 | 0.33 | 12,108 | 62.05% | 15,714 | 56.12% | 28,610 | 50.80% | 13,863 | 29.30% | 33,070 | 14.34% | 24,760 | 40.83% | 10,523 | 17.37% | 54,545 | 8.45% | 25,099 | 14.27% | 8,280 | 6.47% |
| 350-550bp#bulk2 | 99.71 | 62.50 | 2,235,371 | 0.33 | 0.33 | 11,160 | 57.19% | 14,390 | 51.39% | 25,993 | 46.15% | 11,970 | 25.30% | 31,151 | 13.51% | 23,701 | 39.09% | 9,359 | 15.44% | 46,886 | 7.27% | 21,711 | 12.34% | 6,934 | 5.42% |
| 350-550bp#bulk3 | 99.65 | 63.93 | 3,179,090 | 0.34 | 0.34 | 13,058 | 66.92% | 17,435 | 62.27% | 30,224 | 53.66% | 14,669 | 31.01% | 34,250 | 14.85% | 25,443 | 41.96% | 11,828 | 19.52% | 60,074 | 9.31% | 27,168 | 15.45% | 8,968 | 7.01% |
| 350-550bp#bulk4 | 99.66 | 58.09 | 2,855,146 | 0.33 | 0.33 | 12,687 | 65.02% | 17,449 | 62.32% | 28,623 | 50.82% | 13,251 | 28.01% | 33,153 | 14.38% | 24,663 | 40.67% | 11,529 | 19.03% | 56,839 | 8.81% | 24,836 | 14.12% | 7,810 | 6.10% |
| 175-550bp#bulk1 (merged) | 99.66 | 66.22 | 5,889,725 | 0.34 | 0.33 | 16,839 | 86.30% | 24,699 | 88.21% | 37,601 | 66.76% | 19,263 | 40.72% | 40,857 | 17.72% | 28,018 | 46.21% | 20,864 | 34.43% | 97,896 | 15.17% | 36,656 | 20.84% | 12,149 | 9.50% |
| 175-550bp#bulk2 (merged) | 99.67 | 69.16 | 5,253,547 | 0.34 | 0.33 | 16,664 | 85.40% | 24,209 | 86.46% | 36,073 | 64.05% | 17,736 | 37.49% | 39,917 | 17.31% | 27,531 | 45.40% | 20,059 | 33.10% | 90,778 | 14.07% | 33,622 | 19.12% | 10,739 | 8.39% |
| 175-550bp#bulk3 (merged) | 99.65 | 70.23 | 6,992,947 | 0.35 | 0.34 | 17,084 | 87.55% | 25,115 | 89.70% | 38,703 | 68.72% | 20,353 | 43.02% | 41,884 | 18.16% | 28,425 | 46.88% | 21,978 | 36.27% | 104,205 | 16.15% | 38,843 | 22.08% | 13,155 | 10.28% |
| 175-550bp#bulk4 (merged) | 99.67 | 65.46 | 5,938,643 | 0.34 | 0.33 | 16,832 | 86.26% | 24,683 | 88.15% | 36,913 | 65.54% | 18,298 | 38.68% | 40,566 | 17.59% | 27,822 | 45.88% | 20,874 | 34.45% | 95,593 | 14.81% | 35,144 | 19.98% | 11,381 | 8.90% |
| 175-750bp#bulk1 | 99.61 | 69.37 | 6,404,975 | 0.33 | 0.32 | 16,936 | 86.79% | 24,812 | 88.61% | 37,859 | 67.22% | 19,636 | 41.51% | 41,107 | 17.83% | 28,099 | 46.34% | 21,081 | 34.79% | 99,422 | 15.41% | 36,955 | 21.01% | 12,283 | 9.60% |
| 175-750bp#bulk2 | 99.70 | 70.94 | 6,414,211 | 0.33 | 0.32 | 16,991 | 87.08% | 24,791 | 88.54% | 38,337 | 68.07% | 20,207 | 42.72% | 41,524 | 18.01% | 28,348 | 46.75% | 21,389 | 35.30% | 100,563 | 15.58% | 38,057 | 21.64% | 13,003 | 10.16% |
| 175-750bp#bulk3 | 99.67 | 72.43 | 7,188,812 | 0.34 | 0.33 | 17,097 | 87.62% | 25,115 | 89.70% | 38,907 | 69.08% | 20,678 | 43.71% | 41,932 | 18.18% | 28,466 | 46.95% | 21,963 | 36.24% | 104,677 | 16.22% | 38,810 | 22.07% | 13,237 | 10.35% |
| 175-750bp#bulk4 | 99.66 | 69.25 | 6,473,515 | 0.33 | 0.32 | 16,986 | 87.05% | 24,832 | 88.69% | 38,334 | 68.06% | 20,135 | 42.56% | 41,510 | 18.00% | 28,385 | 46.81% | 21,517 | 35.51% | 101,070 | 15.66% | 38,173 | 21.70% | 13,082 | 10.23% |
